# DCX Enhances Glioblastoma Metabolism Through Synergistic Regulation of Glutamine Synthesis and Metabolism-Related Genes for Cellular Homeostasis

**DOI:** 10.1101/2025.06.16.660014

**Authors:** Abiola Abdulrahman Ayanlaja, Xiaoliang Hong, Bo Cheng, Han Zhou, Michael Chang, Kouminin Kanwore, Adebukunola Idayat Adesanya, Motunrayo Monsurat Ayanlaja, Qaudri Akorede Raji, Nadeem Iqra, Piniel Alphayo-Kambey, Chuanxi Tang, Jiaqi Dong, Baole Zhang

## Abstract

Gliomas are the most common primary intracranial tumors, comprising 81% of malignant brain tumors, and currently lack effective therapies. Recent advances in molecular biology have shown that cancer cells exploit microtubule-associated proteins (MAPs) under stress to activate various signaling pathways. This study investigates the role of Doublecortin (DCX) in glioma metabolism and its impact on tumor proliferation. In this study, CRISPR-engineered glioma models with DCX overexpression or knockdown were analyzed using integrated genomic, transcriptomic, and metabolomic approaches. Metabolic activity was assessed via RNA sequencing, Seahorse assays, and targeted mass spectrometry. Pharmacological inhibition of key pathways validated functional dependencies. We demonstrate that gliomas enriched with DCX exhibit elevated glycolytic activity while also relying on cellular respiration and oxidative phosphorylation (OXPHOS) for energy to support the abnormal proliferation of glioma cells. Upon integrative analysis of enriched genes and proteins, we observed genetic and metabolome-level signatures associated with differences in central carbon and energy metabolism in CRISPR-modified glioma cells expressing high DCX. Whole-genome transcriptome analysis revealed enriched metabolic entities promoting hydrolysis of glutamine and glutaminolysis in glioma cells and inhibition of selected differentially enriched genes with small molecule inhibitors abrogated metabolic enrichment and resulted in reduced energy levels and protein translation required for aberrant growth. Finally, we establish that DCX stimulates glutaminolysis to regulate homeostasis for energy supplements in glioma cells. Targeting DCX-mediated metabolic pathways may provide a novel therapeutic approach for glioblastoma, highlighting the potential for innovative treatments in this challenging disease.

## Introduction

Glioblastoma (GBM, World Health Organization grade IV glioma) is the most lethal and commonest form of malignant brain tumor in adults (median survival < 14 months), and it is a very difficult disease to treat ^1^. GBM is characterized by diffused infiltration, drug resistance, metabolic exorbitance, and aberrant proliferation due to the heterogeneity of the GBM population in a tumor mass and the uniqueness of the GBM growth network ^2^. Although GBM cells rely largely on glycolysis to produce ATP even in the presence of oxygen, there is a higher energy demand in ATP for their uncontrollable proliferation. Advances in molecular biology suggest that GBM’s resistance to standard chemotherapy relies on increased metabolic activities via mitochondrial oxidative phosphorylation (OXPHOS) fueling major anti-apoptotic signaling and resistance to therapy ^3–7^. However, a fuller mechanistic understanding of the metabolic processes and the effectors driving it in GBM is still elusive.

The traditional functions of microtubule (MT)-associated proteins (MAPs) have evolved into metabolic accessories ^8^. The MT undergoes conformational changes in response to oxygen deprivation, which may arise from uncontrolled proliferation, and can instigate aberrant expression of MAPs as a response signal. Hence, the cytoskeleton orchestrates cellular signaling via MAPs in response to specific stressors ^9^. One of such MAPs is Doublecortin (DCX), a nervous system-specific MAP crucial for the formation and the proliferation of progenitor cells during neurogenesis ^10–13^. Due to its crucial roles in adult neurogenesis, DCX has underlying roles in oncogenesis and glioma development. It is strongly expressed at the invasive front of gliomas regardless of the tissue grade or type ^14,15^, in patients-derived primary or recurrent GBM ^16^, or cell lines ^17^, and it contributes to the growth and development of GBM cells and enhances the invasive capabilities of GBM cells in mice brain ^15^. DCX also promotes prostate cancer aggressiveness and recurrence. DCX-positive neural progenitor cells originating from the central nervous system (CNS) infiltrate prostate tumors and promote tumor growth metastasis indicating a unique crosstalk between neurogenesis and the initiation of non-CNS tumors ^18^. Since the detection of a comprehensible relationship between DCX expression and cancer malignancy ^14–20^, unraveling the nexus between high expression of DCX, its functions in GBM progression and anti-apoptotic ability, or drug resistance is imperative.

Here, we reveal that DCX promotes amino acid metabolism and regulates cellular homeostasis to stimulate cellular respiration and OXPHOS for energy supplements in GBM cells.

## Materials and Methods

### Cell Culture

Human astrocytes-hippocampal cells (HA-h, Sciencell), human glioma cell lines (U251, LN229, and U87), and rat C6 astroglioma cells were cultured in a humidified incubator at 37°C with 5% CO₂ as previously described^15^. Cells were maintained in DMEM high-glucose media (Hyclone, SH30243.01), DMEM low-glucose media (Hyclone, SH30284.01), or glutamine-deficient DMEM media (Gibco, 11960-044), all supplemented with 10% fetal bovine serum (FBS) and 1% penicillin-streptomycin. For glucose-free conditions, glucose-deficient DMEM media (Gibco, 11966-025) was supplemented with 10% FBS, 1% penicillin-streptomycin, 10 mM D-(+)-Galactose (Sigma, G0750), 1 mM Sodium Pyruvate (Gibco, 11360-070), 2mM glutamine (Gibco, A2916801) and 5mM HEPES (Beyotime, ST523) to create galactose media, which served as an alternative energy source in the absence of glucose. For hypoxia experiments, cells were placed in a hypoxia chamber or incubator maintained at 0.3%-1% O₂, 5% CO₂, and 94% N₂ at 37°C, with media pre-equilibrated to the hypoxic conditions prior to use. Cells were passaged at 80-90% confluency using 0.25% trypsin-EDTA (Gibco, 25200-056) and reseeded at appropriate densities, with media replaced every 2-3 days to ensure optimal growth conditions.

### Seahorse Extracellular Flux Analysis

Extracellular flux analysis was conducted using the Seahorse XF24 platform (Agilent, 102601-100) to evaluate metabolic parameters in glioma cells. Cells were uniformly seeded into XF24 plates and maintained under varying culture conditions for 24–48 hours. Metabolic measurements were performed using the Seahorse XF24 analyzer and Wave 2.6.1 software (Agilent). To assess substrate utilization and metabolic flexibility, U251 cells were subjected to stress tests using all three substrate test kits. To eliminate bias in substrate uptake measurements, cells were cultured in glutamine-deficient media (DMEM No-Glutamine), enabling dual inhibition of extracellular and intracellular glutamine through media composition and BPTES treatment. Additionally, cells were cultured in low-glucose and no-glucose (galactose-based) conditions to evaluate glucose dependency. For the glutamine stress test, injection ports of the XFe24 Seahorse cartridge were loaded with BPTES, oligomycin, FCCP, and rotenone plus antimycin A. In glucose/pyruvate stress tests, UK5099 replaced BPTES. For long-chain fatty acid (LCFA) oxidative stress tests, etomoxir was used instead of BPTES or UK5099. These protocols allowed for comprehensive assessment of mitochondrial respiration (OCR) and glycolytic flux (ECAR) under varying metabolic stresses, providing insights into substrate preferences and metabolic vulnerabilities in glioma cells.

### Amino Acid Composition Analysis

1×10⁷ cells (DCX-OE and U251 control) were collected for amino acid profiling by Shanghai Biotree Biotech Co. Ltd. (China) using ultra-high-performance liquid chromatography (UHPLC) on an Agilent 1290 Infinity II series system (Agilent Technologies). Separation with a Waters ACQUITY UPLC BEH Amide column (100 mm × 2.1 mm, 1.7 μm). The mobile phases consisted of (A) 1% formic acid in water and (B) 1% formic acid in acetonitrile. The column temperature was maintained at 35°C, and the auto-sampler was set to 4°C with an injection volume of 1 μL.

Detection was carried out using an Agilent 6460 triple quadrupole mass spectrometer equipped with an AJS electrospray ionization (AJS-ESI) interface. Ion source parameters included a capillary voltage of +4,000 V (positive mode) or −3,500 V (negative mode), nozzle voltage of +500 V (positive mode) or −500 V (negative mode), nitrogen gas temperature of 300°C, gas flow of 5 L/min, sheath gas temperature of 250°C, sheath gas flow of 11 L/min, and nebulizer pressure of 45 psi. Multiple reaction monitoring (MRM) parameters were optimized for each analyte via flow injection analysis of standard solutions. The most sensitive Q1/Q3 transition was selected as the quantifier for quantification, while additional transitions served as qualifiers for analyte confirmation. Data acquisition and processing were performed using Agilent MassHunter Work Station Software (B.08.00, Agilent Technologies).

### Metabolite Profiling Analysis

Glioma cells (DCX-OE and U251 control; 1×10⁷) were collected for metabolomics assay. Metabolomic profiling was performed at Shanghai Biotree Biotech Co. Ltd. (China). Briefly, 500 μl of precooled MeOH/H2O (3:1, v/v) was added to each sample, followed by vortexing for 30 seconds. Samples were flash-frozen in dry ice and subjected to three freeze-thaw cycles in liquid nitrogen. After vortexing again for 30 seconds, samples were sonicated for 15 minutes in an ice-water bath. Subsequently, samples were incubated at -40 °C for 1 hour and centrifuged at 12,000 rpm (RCF = 13,800 g, radius = 8.6 cm) for 15 minutes at 4 °C. A 400 μl aliquot of the clear supernatant was collected and dried by centrifugation. The residue was reconstituted in 200 μl of ultrapure water, vortexed, and filtered using a centrifuge tube filter before being transferred to injection vials for HPLC-MS/MS analysis. HPLC separation was performed using a Thermo Scientific Dionex ICS-6000 system equipped with Dionex IonPac AS11-HC (2 × 250 mm) and AG11-HC (2 × 50 mm) columns. Metabolite detection was carried out using an AB SCIEX 6500 QTRAP+ triple quadrupole mass spectrometer with an electrospray ionization (ESI) interface operating in multiple reaction monitoring (MRM) mode. Data acquisition and processing were conducted using AB SCIEX Analyst Work Station Software (1.6.3), MultiQuant 3.0.3, and Chromeleon 7. The final dataset, including compound names, sample identifiers, and concentrations, was imported into SIMCA 16.0.2 (Sartorius Stedim Data Analytics AB, Umea, Sweden) for multivariate analysis. Data were scaled and log-transformed to reduce noise and variance. Statistical analysis and visualization were performed using R packages, including tidyverse, dplyr, magrittr, ggplot2, and ggrepel for volcano plot construction, circlize and reshape2 for chord diagrams, and pheatmap for heatmap generation.

### Promoter Activity Validation

The promoter activity of the human ASNS gene was validated using a dual-luciferase reporter assay system provided by Hanbio Biotechnology Co., Ltd. (Shanghai, China). The ASNS promoter region (2000 bp upstream of the transcription start site) was cloned into the pGL3-basic vector upstream of the Firefly luciferase (F-Luc) gene. The transcription factor ATF5 was cloned into the pcDNA3.1 vector for co-expression. Mutant versions of the ASNS promoter, with mutated predicted ATF5 binding sites, were also constructed. 293T cells were transfected with the following combinations: (1) ASNS promoter-luc-NC+ATF5-mut1, (2) ASNS promoter-luc-NC+ATF5-mut2, (3) ASNS promoter-luc+ATF5, (4) ASNS promoter-luc+ATF5-OE. These combinations were repeated in DCX-OE cells. Transfections were performed using LipoFiter™ transfection reagent (Hanbio Biotechnology) according to the manufacturer’s protocol. Cells were harvested 48 hours post-transfection, and luciferase activity was measured using the Dual-Luciferase® Reporter Assay System (Promega). Firefly luciferase activity was normalized to Renilla luciferase activity for each sample.

### In Vivo Mouse Studies

Male BALB/c nude mice (immunocompromised) or immunocompetent BALB/c mice, aged 6–8 weeks, were obtained from the Animal Care Center of Xuzhou Medical University. Eight mice were allocated to each experimental cohort (control, DCX-OE, and DCX-KD groups). Mice were housed and handled in accordance with the National Institutes of Health Guide for the Care and Use of Laboratory Animals. Mice were monitored daily by an independent, blinded evaluator for tumor progression and neurological symptoms (e.g., lethargy, ataxia, or weight loss >20%). Animals exhibiting severe neurological impairment were humanely euthanized via CO₂ inhalation followed by cervical dislocation. All animal experiments were conducted under protocols (202212S001) approved by the Animal Care Committee of Xuzhou Medical University as previously described^15^. Kaplan-Meier survival curves were generated, and statistical significance between groups was determined using the log-rank test in GraphPad Prism v10.2 software (GraphPad Software, Inc.).

### siRNA Transfection

Stealth siRNA targeting asparagine synthetase (ASNS) was purchased from Invitrogen. According to manufacturer’s protocol, siRNA was resuspended in DEPC-treated H₂O to prepare a 20 μM stock solution, which reconstituted the buffer to 10 mM Tris-HCl (pH 8.0), 20 mM NaCl, and 1 mM EDTA. 0.5–5 pmol of siRNA was transfected by adding 100 μL of transfection mix to 500 μL of medium, achieving a final concentration of 0.8–8 nM in a 24-well plate. For scaling up, the siRNA amount was increased proportionally to maintain the desired concentration.

### Determination of ATP concentration

100 μL of ATP detection working solution was added each detection well of a 96-well detection plate. To expel background ATP, the solution was left in the detection plate for 3-5 minutes at room temperature. Then, 20 μL of sample or standard was added to the test hole and mixed appropriately without creating bubbles. Within 2 seconds, RLU value Or CPM was detected in Biotek’s Synergy™ 2 Multi-Mode Microplate Reader. ATP concentration was then calculated according to the standard curve.

### NAD /NADH detection

Cellular NAD+/NADH levels were quantified using Beyotime’s NAD^+^/NADH assay kit (WST-8 method). Approximately 1×10⁶ cells were seeded in a 6-well plate, lysed with 200 μl of NAD^+^/NADH extraction buffer, and centrifuged at 12,000 × g for 10 minutes at 4°C. A 10 mM NADH standard was prepared and serially diluted to generate a concentration gradient (0–10 μM). For the assay, 20 μl of each sample or standard was added to a 96-well plate, followed by 90 μl of alcohol dehydrogenase working solution. After incubation at 37°C for 10 minutes in the dark, 10 μl of color developing solution was added, and the plate was incubated for an additional 30 minutes at 37°C. The absorbance of the orange-yellow formazan product was measured at 450 nm using a Biotek Synergy™ 2 Multi-Mode Microplate Reader. A standard curve was generated by plotting NADH concentration against absorbance. The NAD+ concentration was calculated as the difference between total NAD (NAD+ + NADH) and NADH levels, and the NAD+/NADH ratio was determined using the formula: NAD+/NADH=([NAD total]−[NADH])/[NADH].

### Lactate Detection

Cellular lactate levels were quantified using BioVision’s Lactate Colorimetric Assay Kit II (#K627). Briefly, Lactate Enzyme Mix and Lactate Substrate Mix were reconstituted in 0.22 mL of Lactate Assay Buffer. For the standard curve, the 100 mM L(+)-Lactate Standard was diluted to 1 mM by adding 10 μL of the standard to 990 μL of Lactate Assay Buffer. Serial dilutions (0–10 μL) were added to a 96-well plate, adjusting the volume to 50 μL/well with Lactate Assay Buffer to generate 0–10 nmol/well of lactate. Cell samples were prepared at 50 μL/well in the same plate. Then, 50 μL reaction mix, consisting of 46 μL Lactate Assay Buffer, 2 μL Lactate Substrate Mix, and 2 μL Lactate Enzyme Mix, was added to each well containing standards or samples. The plate was incubated at room temperature for 30 minutes, and absorbance was measured at 450 nm using a Biotek Synergy™ 2 Multi-Mode Microplate Reader. Lactate concentration was calculated using the formula: C=LaSvl1(nmol/μL or mM), where La is the lactate amount (nmol) derived from the standard curve, and Sv is the sample volume (μL) added to the well.

### Glutathione (GSH) Detection

Cellular glutathione (GSH) levels were measured using Abcam’s Glutathione Colorimetric Detection Kit. A 5% (w/v) 5-sulfo-salicylic acid dihydrate (SSA) solution was prepared by dissolving 1 g of SSA in 20 mL of ddH₂O. The sample diluent buffer was prepared by mixing 5 mL of 5% SSA with 20 mL of assay buffer, followed by thorough vortexing and pH adjustment to >6. Cell pellets were washed with ice-cold PBS and resuspended in ice-cold 5% SSA at a concentration of 1–40 × 10⁶ cells/mL. Cells were lysed by sonication and incubated for 10 minutes at 4°C. Lysates were centrifuged at 14,000 rpm for 10 minutes at 4°C, and the supernatant was collected for analysis. GSH levels were determined by colorimetric measurement using a Biotek Synergy™ 2 Multi-Mode Microplate Reader by measuring the rate of color development at 405 nm.

### Protein Extraction, Immunoblot and Immunoprecipitation Assay (IP)

Whole cell lysates were extracted using cell lysis buffer (Beyotime), supplemented with protease inhibitor coctails and PMSF. Protein concentration was measured using BCA protein assay kit (ThermoFisher). Immunoblot and co-immunoprecipitation were performed as previously described ^15^.

### RT-qPCR

Total RNA extraction and RT-qPCR were carried out as previously described ^21^. Data was normalized to stable reference genes (β-actin) using the ΔCt method, and relative gene expression was calculated as 2^(-ΔΔCt). Statistical comparisons were performed using unpaired t-tests (two groups) or one-way ANOVA with Tukey’s post-hoc test (three or more groups), with non-parametric tests (Mann-Whitney U) applied for non-normal data. Data are presented as mean ± SEM, with significance defined as p < 0.05 and adjusted for multiple testing using the Benjamini-Hochberg method.

### Transmission Electron Microscopy

Transmission electron microscopy (TEM) sample preparation was performed as previously described ^22^. Briefly, samples were chemically fixed using aldehydes to cross-link proteins, followed by osmium post-fixation to stabilize macromolecular structures. Dehydration was carried out using a graded acetone series (40%, 70%, 90%, and 100% acetone, 10–20 minutes each, repeated three times). For infiltration, acetone was replaced with an epoxy resin (Epon) intermediary solvent. Samples were incubated in 1:1 Epon–acetone for 30 minutes, followed by 3:1 Epon–acetone for 45–60 minutes, and finally in 100% Epon twice for 1 hour. Embedding was performed using pre-dried molds, and samples were polymerized at 70°C in a vacuum oven for 48 hours. Sections of 50–500 nm thickness were obtained using ultramicrotomy, mounted on 3 mm copper grids pre-coated with a 0.1 μm carbon film, and double-stained with 1% uranyl acetate and alkaline lead citrate. Micrographs were acquired as high-resolution black-and-white images.

### RNA Sequencing

RNA sequencing was conducted and analyzed by Shanghai Biotechnology Corporation (China). Total RNA was isolated from 1×10⁶ cardiac macrophages using the RNeasy Micro Kit (74004, Qiagen) according to the manufacturer’s protocol. RNA quality was assessed, and qualified samples were used for library preparation and sequencing. Differentially expressed genes (DEGs) were identified and filtered using the DEGseq algorithm in R, with p-values adjusted via Benjamini & Hochberg’s False Discovery Rate (FDR) correction method (adjusted p < 0.05). Gene set enrichment analysis (GSEA) was conducted using the clusterProfiler v4.8.3 package in R. Genes were pre-ranked based on the signed log2 fold change multiplied by -log10(p-value), as calculated by DESeq2. The MsigDBR v7.5.1 package was employed to query gene sets from the Hallmark (H), KEGG (CP:KEGG ), and chemical and genetic perturbations (CGP) collections ^23,24^. Additionally, hypergeometric enrichment analysis was performed using the clusterProfiler package. Pathway annotation was carried out using Hallmark gene sets and oncogenic signature gene sets from the Molecular Signatures Database ^25^.

### Statistical Analyses

To identify differentially expressed genes (DEGs), the Wald test was used in combination with DESeq, a tool for RNA-seq data analysis. Results were visualized using volcano plots, with DEGs defined by a fold change of 2 or greater (either up or down, indicated by vertical dashed lines) and a Benjamini-Hochberg adjusted p-value of ≤ 0.05 (indicated by a horizontal dashed line). Transcriptomic signatures across cell lines under different treatment conditions were analyzed using Student’s t-test. For gene set enrichment analysis (GSEA), p-values were adjusted using the Benjamini-Hochberg method as the default.

## Results

### DCX Promotes Glioblastoma Proliferation and Tumor Progression

Previous findings by us and others showed that elevated DCX expression promotes glioma growth^15^. To further investigate the biological significance of DCX in cancer progression, we generated stable CRISPR/Cas9-based gain-of-function and knockdown models in human glioma cell lines (U251-MG, LN-229, and U87-MG) and rat-derived C6 glioma lines, as previously described^15^. For DCX knockdown (DCX-KD), Sanger sequencing confirmed INDELs and polyclonal pools with expected cleavage patterns were verified using the CRISPR Genomic Cleavage Detection Kit (abm G932).

To explore the downstream effects of DCX modulation, we analyzed signaling changes and differential gene expression in U251-control lines versus U251 cells overexpressing DCX (DCX-OE) using bulk RNA sequencing (RNAseq) (Figure S1A, B). Geneset enrichment analysis (GSEA) identified significant enrichment of key Hallmark cell proliferation-related gene sets^26^, including HALLMARK_G2M_CHECKPOINT (Figure 1A), HALLMARK_MYC_TARGETS_V1 (Figure 1B), and HALLMARK_E2F_TARGETS (Figure 1C).

**Figure 1.**
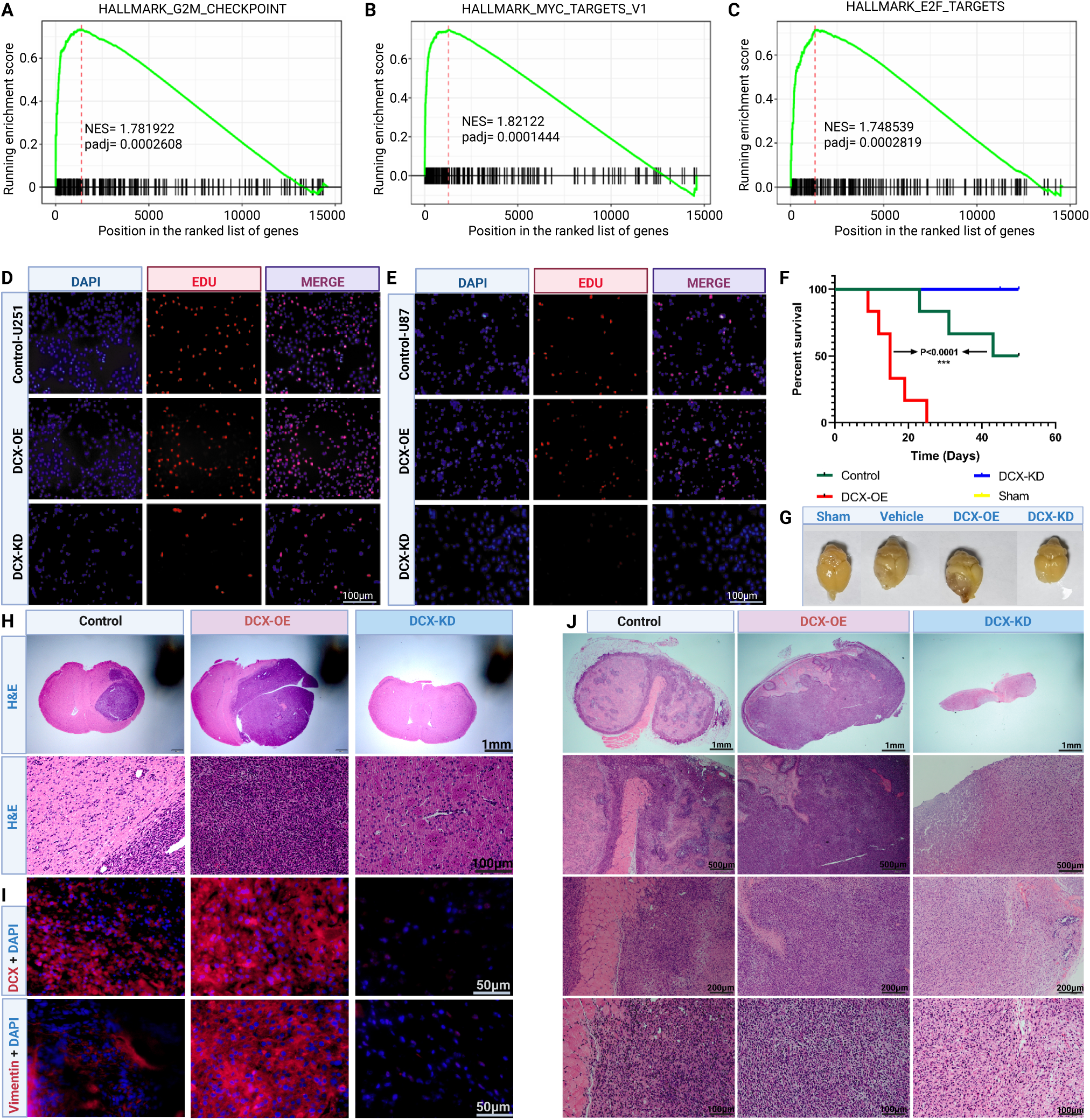
DCX Promotes Glioblastoma Proliferation and Tumor Progression: Genset enrichment analysis of DCX-OE – control for gene sets of interest including A) HALLMARK_G2M_CHECKPOINT, B) HALLMARK_MYC_TARGETS_V1 and C) HALLMARK_E2F_TARGETS. D) Representative fluorescence images of EdU incorporation in U251-MG and E) C6 glioma cells. Blue = nuclei staining for DAPI, red = EdU positive staining. Magnification, 100µm. F) Kaplan-Meier survival curves of BALB/c nude mice implanted intracranially with U251-MG cells. G) Representative whole brain images from each cohort (control, DCX-OE, and DCX-KD) at the study endpoint. H) Hematoxylin and eosin (H&E)-stained sections from mouse brains magnification, 1mm. Lower panels magnified field, 100um. I) Immunofluorescence staining of brain sections showing expression of DCX and vimentin in DCX-OE tumors compared to controls and DCX-KD tumors at 50 µm magnification. J) Histology of subcutaneous tumors in immune-competent BALB/c mice implanted with C6 glioma cells in DCX-OE and DCX-KD tumors compared with control. Magnification, 1mm, 500µm, 200µm, 100µm.

We next assessed the role of DCX on cell cycle dynamics through flow cytometry in U251 and U87-MG lines. DCX-OE cells exhibited a greater proportion of cells in the S-phase compared to controls and DCX-KD cells (Figure S1C). This was supported by EdU incorporation assays (Figures 1D, E), which revealed a significant increase in EdU-positive cells in DCX-OE U251-MG and U87-MG cells, while DCX-KD cells showed markedly reduced EdU incorporation, indicating diminished proliferation.

The *in vivo* effects of DCX on GBM growth were assessed using orthotopic intracranial and heterotopic subcutaneous xenograft models. In the intracranial model, U251-MG cells were implanted into immune-compromised BALB/c nude mice, whereas the subcutaneous model involved the implantation of C6 cells into flanks of immune-competent BALB/c mice. Mice implanted with DCX-OE cells demonstrated significantly shorter survival times compared to controls, while DCX-KD cohorts survived until the study endpoint (Figure 1F). Histological analysis further validated these findings. Tumor sizes, visualized through whole-brain imaging, and hematoxylin and eosin (H&E) staining, were markedly larger in DCX-OE mice, with features consistent with aggressive GBM histopathology (Figure 1G, H). Immunofluorescence staining revealed elevated expression of DCX and vimentin DCX-OE tumors. Conversely, control tumors were smaller with reduced DCX and vimentin expression. DCX-KD mice exhibited minimal or no detectable DCX expression, lower vimentin levels, and significantly smaller tumor sizes (Figure 1I). Similar trends were observed in the subcutaneous xenograft model (Figure 1J). C6 cells overexpressing DCX formed larger flank tumors, while DCX knockdown significantly suppressed tumor growth.

### DCX-Induced Metabolic Reprogramming Fuels Glioma Cell Proliferation

To assess transcriptional changes in DCX-OE cells compared to controls, we identified the top 10 upregulated and downregulated genes. Upregulated genes were associated with protein translation, glutamine metabolism, amino acid exchange, and cellular stress response (Figure S2A). Moreover, GO analysis indicated the enrichment of several biological processes including metabolism, response to stimuli, and proliferation (Figure S2B), suggesting global metabolic reprogramming. GSEA revealed reduced glucose metabolic process in DCX-OE cells when compared with control cells (Figure 2A), prompting further investigation of metabolic enzyme activity.

**Figure 2.**
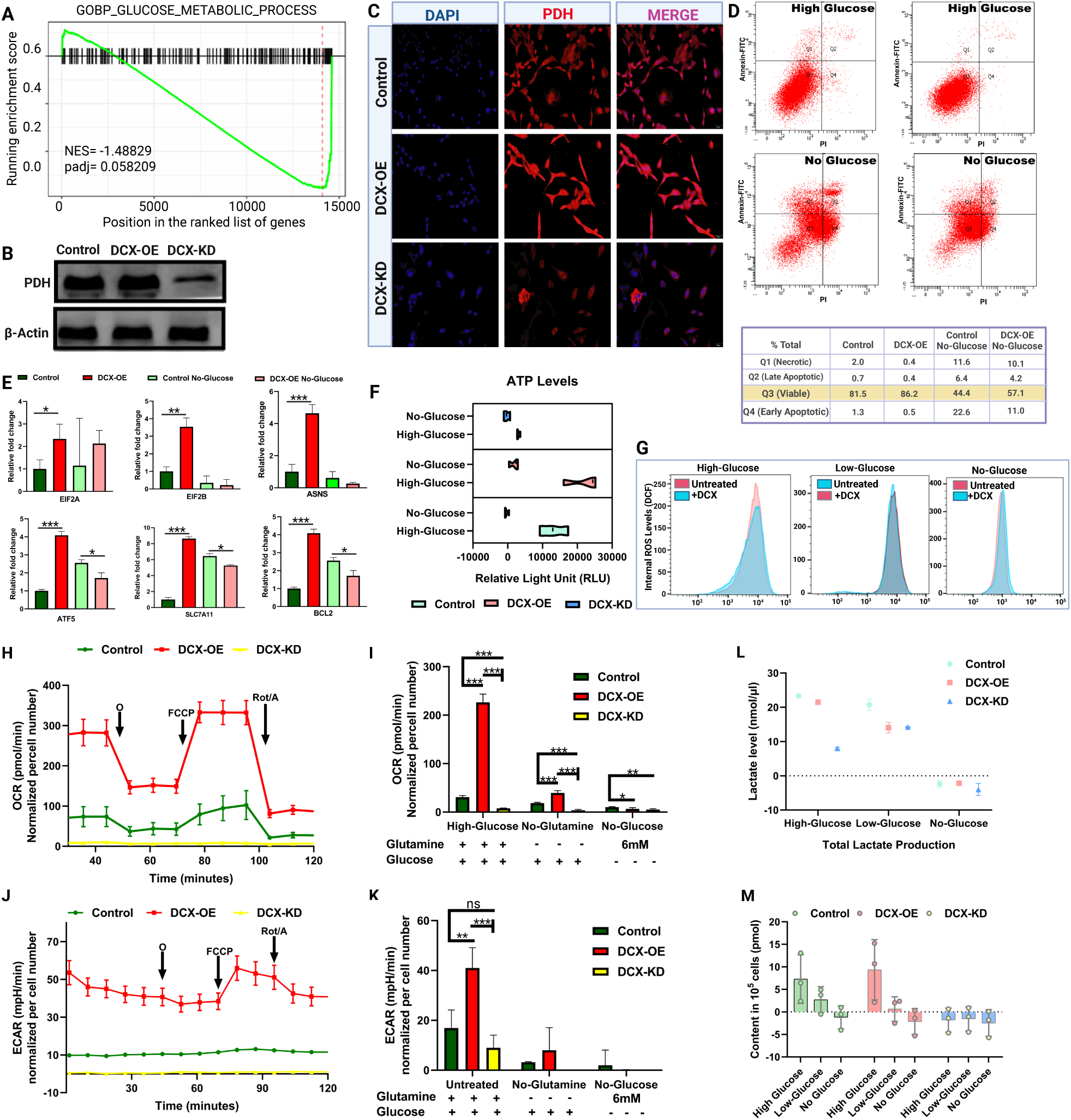
DCX promotes glioma proliferation via metabolic reprogramming in glioma cells: A. Genset enrichment analysis of DCX-OE vs control showing GOBP_GLUCOSE_METABOLIC_PROCESS. B) Immunoblots of lysates from control, DCX-OE, and DCX-KD cells showing pyruvate dehydrogenase (PDH) and actin. C) Fluorescence imaging of PDH (red), and DAPI (blue) in U251-Control, DCX-OE, and DCX-KD cells, 50µm. D) Flow cytometry analysis of cells cultured in high-glucose (normal) and glucose-deficient culture conditions (galactose media). E) Relative EIF2A, EIF2B, ASNS, ATF5, SLC7A11, and BCL2 mRNA expression levels in control and DCX-OE cells cultured in normal and galactose media culture conditions. Results are presented as mean ± SEM. *P < 0.05, **P < 0.01, and ***P < 0.001. F) Fluorometric detection of ATP levels in control, DCX-OE, and DCX-KD cells. G) Flow cytometry analysis of internal ROS levels of cells stained with DCF dichlorodihydrofluorescein (DCF), wild-type GBM cells were cultured in normal, low-glucose, and galactose media culture conditions and treated with or without recombinant DCX protein. H) Seahorse metabolic assay measuring oxygen consumption rate (OCR) in Control-U251, DCX-OE, and DCX-KD cells. O, oligomycin; F, FCCP (carbonyl cyanide 4-[trifluoromethoxy] phenylhydrazone); A&R antimycin A and rotenone. I) Analysis of maximum OCR levels in normal culture conditions, no-glutamine, and galactose media. Data are mean + SD (n=3-4). Student’s t-test. *P < 0.05, **P < 0.01, ***P < 0.001. J. ECAR levels in normal culture condition, no-glutamine, low-glucose, and galactose media, and K) analysis of maximum ECAR. Data are mean + SD (n=3-4). Student’s t-test. *P < 0.05, **P < 0.01, ***P < 0.001. L. Fluorometric detection of lactate levels in control, DCX-OE, and DCX-KD cells. M. Fluorometric detection of NAD/NADH levels in control, DCX-OE, and DCX-KD cells.

DCX overexpression stimulated glucose recycling, evidenced by increased mRNA levels of Glucose-6-phosphatase (G6PC) (Figure S2C), a key enzyme in glucose homeostasis^27^. Immunoblot (Figure 2B) and immunofluorescence imaging (Figure 2C) demonstrated enriched pyruvate dehydrogenase (PDH), indicating enhanced glycolytic flux. While DCX upregulated glycolytic enzymes, lower mRNA levels of *PFKFB* suggested limited glycolytic energy output (Figure S2D). Since PFKFB3 inhibition is known to suppress glycolysis and tumor growth^28^, these findings imply that optimal energy requirements for high-DCX-induced proliferation extend beyond glycolysis.

To investigate reliance on alternative energy sources, U251-MG cells were cultured in low-glucose or glucose-deficient media supplemented with galactose, changing the carbon source and thereby reducing glycolytic flux, forcing OXPHOS. DCX-OE cells showed reduced apoptosis and necrosis in these conditions (Figure 2D), indicating alternative metabolic adaptations. However, mRNA expression of top DEGs, the EIF2 complex, and *BCL2* declined in galactose media, while pro-apoptotic BAX levels remained low (Figure 2E, S2E), suggesting glucose is critical for maintaining transcriptional activity of key genes.

Cancer cells require rapid ATP production and tightly regulated ROS metabolism for redox balance and survival^29^. DCX-OE cells maintained consistent ATP and ROS levels across glucose-replete and galactose conditions (Figure 2F, G). In hypoxic conditions, DCX-OE cells exhibited increased growth (Figure S2F), and upregulation of hypoxia-related gene sets (Figure S2G), highlighting metabolic reprogramming driven by DCX to sustain proliferation.

To further explore DCX’s role in glucose-pyruvate oxidation, we conducted Seahorse assays to measure oxygen consumption rate (OCR) and extracellular acidification rate (ECAR). DCX overexpression enhanced OCR under normal conditions, with OCR further reduced in glutamine-deficient media but remaining higher than in control (Figure 2H, I), indicating high glucose demand in glutamine-deficient-culture conditions. The dramatic increase in OCR in DCX-OE cells suggests a shift from glycolytic reliance in control cells to a more balanced energy metabolism that enhances oxidative phosphorylation (OXPHOS). To understand the maximal substrate uptake without bias, we cultured cells in galactose medium, thereby establishing double inhibition of extracellular and intracellular glucose in culture media before subjecting cells to UK5099 for glucose/pyruvate oxidation stress test. In galactose media, OCR dropped sharply, and high DCX expression failed to rescue it (Figure 2I). ECAR analysis revealed increased glycolytic flux in normal media, reduced ECAR in glutamine-deficient conditions, and depleted ECAR in galactose media (Figure 2J, K). DCX-OE cells lactate production was significantly depleted by glucose withdrawal, indicating deregulation of the Warburg effect (Figure 2L). Additionally, DCX enhanced total NAD/NADH levels in a glucose-dependent manner (Figure 2M). These results demonstrate that DCX promotes aerobic glycolysis and metabolic reprogramming in GBM cells, supporting cell proliferation through enhanced glycolytic and oxidative activity.

### DCX promotes amino acid synthesis via glutamine metabolism

To investigate the role of DCX in reprogrammed metabolism in glioma cells, particularly under conditions where glycolysis alone may not meet energy demands, we examined its impact on glutamine metabolism. DCX overexpression significantly upregulated *glutaminase* (*GLS2*) mRNA expression (Figure 3A), indicating rapid glutaminolysis^30^. In cancer cells, ROS levels are tightly regulated to balance oxidative stress and maintain redox homeostasis, ensuring survival and proliferation under metabolic stress. The detrimental impact of ROS on cellular components and signaling pathways can foster conditions that promote conducive environment for cancer cells^31^. In this study, we used dihydroethidium (DHE) staining to quantify intracellular ROS levels in DCX-overexpressing (DCX-OE) glioblastoma cells. Elevated intracellular ROS in DCX-OE cells, detected via Dihydroethidium (DHE) staining was sustained in both normal and glutamine-deficient conditions (Figure 3B). However, high transcription of *TIGAR* would protect DCX-OE from ROS-induced apoptosis (Figure 3C). Immunoblots showed that DCX-OE cells maintained high protein expression of top DEGs including *ASNS* and *xCT* (*SLC7A11*), key regulators of glutamine hydrolysis and transport^32,33^, even under glutamine-deficient conditions (Figure 3D). Glutamine withdrawal led to the complete depletion of DCX protein in DCX-KD cells, correlating with inactivation of the AKT/mTOR signaling and its downstream effectors (Figure 3D).

**Figure 3.**
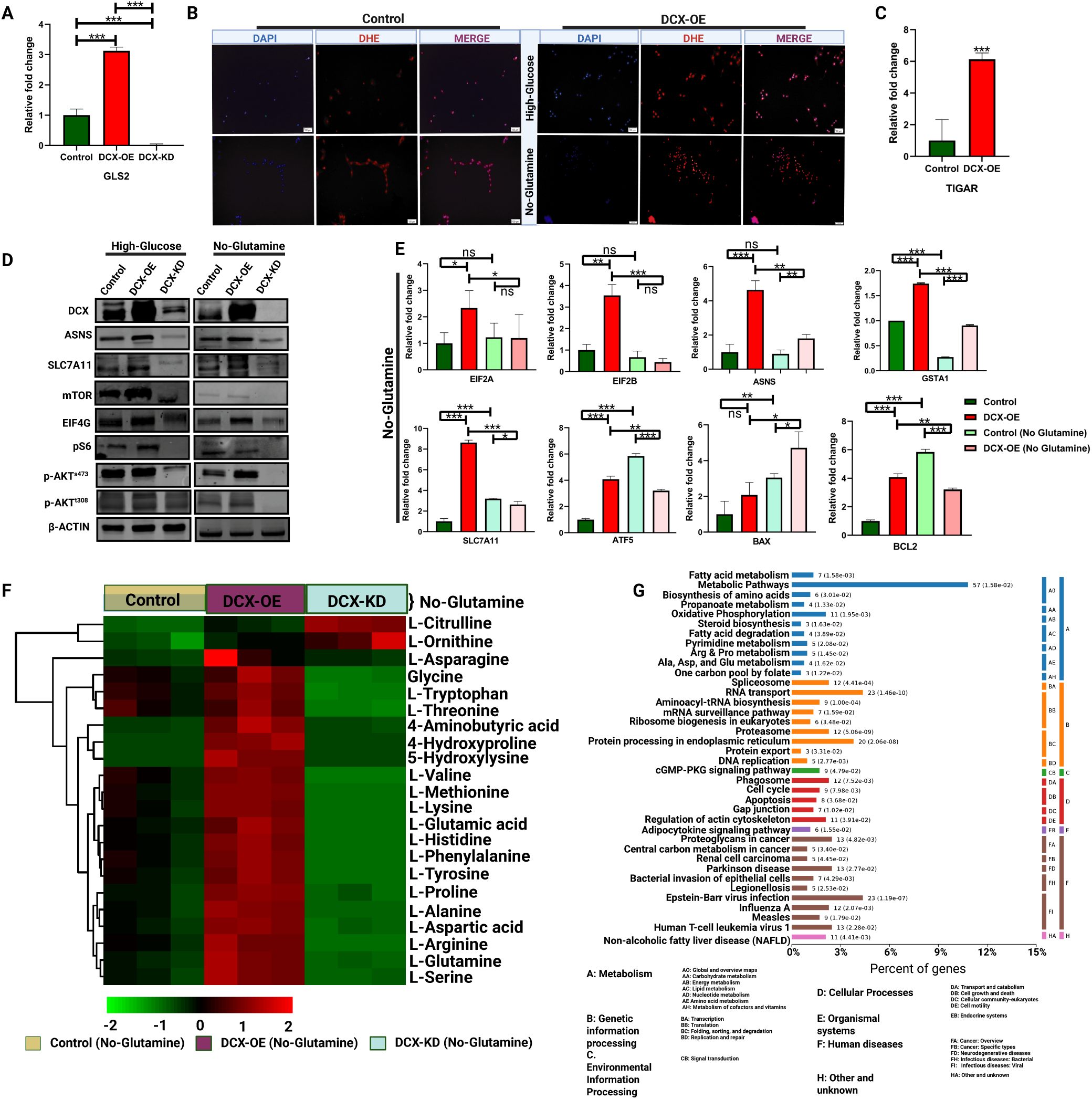
DCX promotes amino acid synthesis via glutamine metabolism | A. Relative fold change of GLS2 mRNA expression in control, DCX-OE, and DCX-KD cells. B. Fluorescence imaging of DHE staining (red) of control and DCX-OE cells cultured in normal and glutamine-deficient conditions. C. Relative fold change of TIGAR mRNA expression in control and DCX-OE cells. D. Immunoblotting of DCX, ASNS, xCT, mTOR, EIF4G, S6K1, pAKT^T308^, pAKT^S473^, and actin in normal and glutamine-deficient condition E. Relative fold change of mRNA levels of EIF2A, EIF2B, ASNS, GSTA1, SLC7A11, ATF5, BAX, and BCL2 in glutamine-deficient culture conditions. F. Intracellular amino acid profiling via UHPLC-MRM-MS/MS in control, DCX-OE, and DCX-KD cells cultured in glutamine-deficient culture condition. G) GO analysis of DCX-interacting proteins identified by mass spectrometry. Results are presented as mean ± SEM. *P < 0.05, **P < 0.01, and ***P < 0.001.

RNAseq revealed enrichment of amino acid metabolic activity related gene sets in DCX-OE cells (Figure S3A). ASNS, a key enzyme in the reversible conversion of glutamate to glutamine, emerged as critical for DCX-mediated metabolic activities. Knockdown of ASNS in U87MG cells using siRNA, or inhibition using cycloheximide (CXH) or L-asparaginase, suppressed top DEGs, *EIF2A, EIF2B, LDHA*, and *BCL2*, and triggered apoptosis via *BAX* upregulation (Figure S3B-D).

Furthermore, glutamine withdrawal also significantly reduced mRNA levels of the EIF2 complex, downregulated top DEGs, and activated pro-apoptotic genes in DCX-OE cells (Figure 3E). However, DCX-OE cells sustained proliferation and S-phase enrichment, likely due to intracellular glutamine retention (Figure S3E. Consistently, DCX-OE cells displayed enhanced EIF4G fluorescence under glutamine-deficient conditions, highlighting active protein synthesis (Figure S3F). Amino acid profiling via UHPLC-MRM-MS/MS demonstrated sustained synthesis of essential amino acids, including alanine, aspartate, and glutamate, in DCX-OE cells, even under glutamine-deficient conditions, though levels were higher in glutamine-replete conditions (Figure 3F, S3G).

Co-immunoprecipitation and mass spectrometry revealed 5,390 differentially expressed proteins in DCX-OE cells, compared to 1,640 in controls. These proteins were enriched in pathways related to amino acid metabolism, oxidative phosphorylation, and biosynthesis of alanine, aspartate, and glutamate (Figure 3G, S3H). KEGG and GO analyses highlighted involvement in glycolysis, TCA cycle, fatty acid metabolism, amino acid transporters and catalytic enzymes, mRNA translation, and protein synthesis, further supporting its role in cellular energy generation and proliferation (Figures S3I). These findings demonstrate that DCX facilitates glioma cell growth and survival by regulating glutamine metabolism, amino acid synthesis, and protein translation, ensuring metabolic adaptability under nutrient stress.

### DCX enhances mitochondrial biogenesis and oxidative phosphorylation in GBM cells

RNAseq analysis further revealed enrichment of mitochondrial matrix activity in DCX-OE cells, a complex containing enzymes of the TCA cycle such as *PDK* (*pyruvate dehydrogenase kinase*), and *GLS2* which catalyzes glutamine hydrolysis (Figure S4A)^34,35^. Subsequent analysis showed that DCX promotes the enrichment of TCA cycle intermediate enzymes, supporting enhanced OXPHOS (Figure 4A). Metabolomics analysis using Ultra-high performance liquid chromatography-mass spectrometry (UHPLC-MS) confirmed elevated TCA cycle intermediates and crucial central carbon metabolites in DCX-OE cells (Figures 4B, S4B). However, glutamine withdrawal resulted in the depletion of energy stores in the form of ATP, ADP, GTP, and TCA intermediates, highlighting DCX-OE cells reliance on glutamine to sustain energy stores and mitochondria activity (Figure S4C, S4D). This indicates that DCX is important for the TCA cycle anaplerosis, and a withdrawal of glutamine affects the energy requirement of DCX-OE cells. However, intracellular glutamine could still maintain homeostasis to regulate oxidative damage (Figure S4D), even though ROS levels were higher in DCX-OE cells after glutamine withdrawal (Figure 3B). In low-glucose growth conditions, enriched TCA cycle was observed in DCX-OE (Figure 4B). Meanwhile, lower energy stores and sustained homeostasis was observed in glucose-deficient growth conditions, indicating glutamine abundance in cells is essential for homeostasis and survival. Predictably, high DCX expression is consistent with higher energy generation in these culture conditions when compared with control in similar conditions. Furthermore, glutamine withdrawal induces depletion of important TCA cycle metabolites and lower energy storage form when compared to low glucose groups with normal glutamine levels (Figure 4B, S4B-S4D).

**Figure 4.**
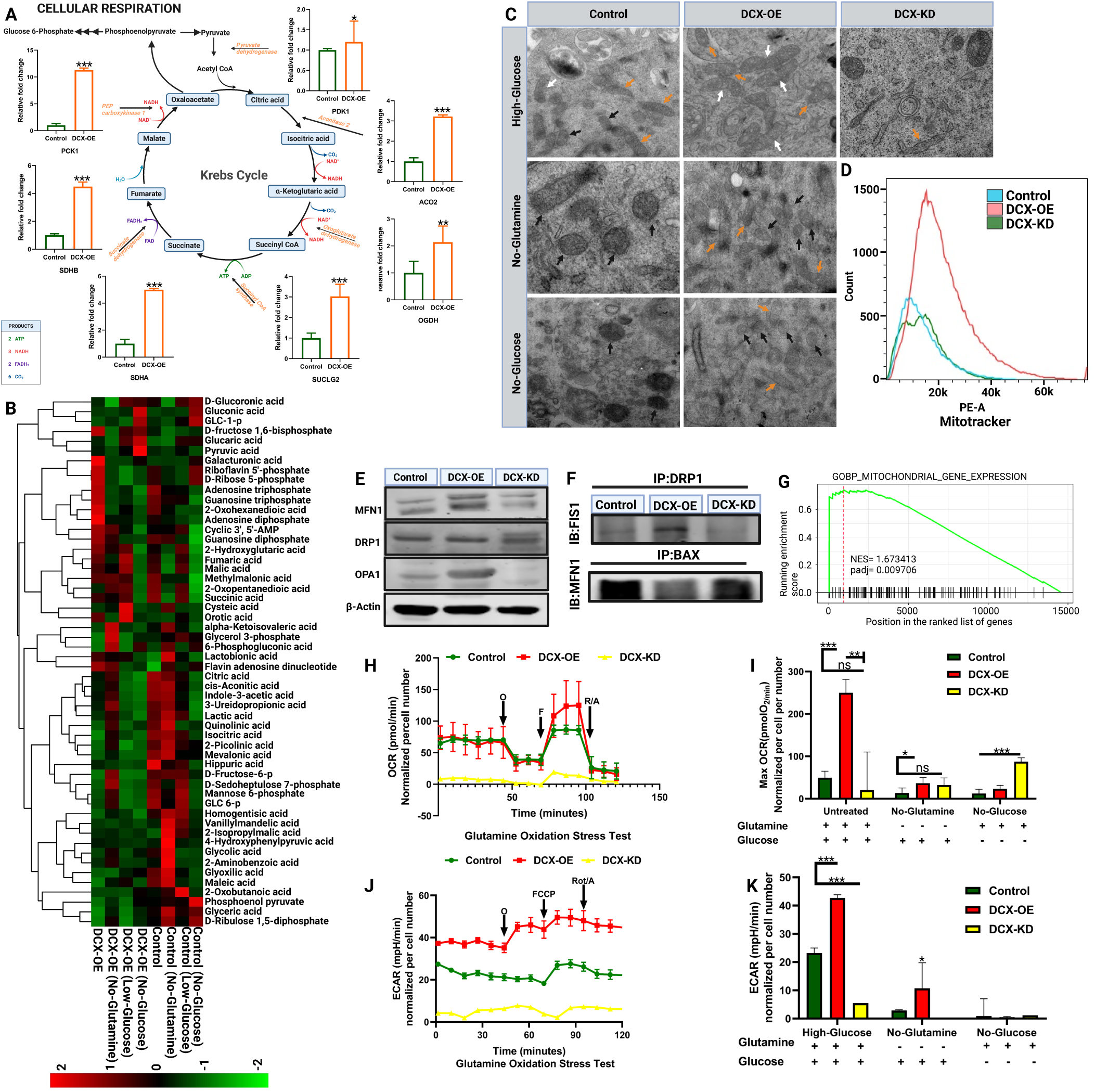
DCX enhances mitochondrial biogenesis and oxidative phosphorylation in GBM cells. A) Relative mRNA expression to actin probing TCA cycle intermediates enzymes in DCX-OE cells compared with control cells. Three independent experiments. Data are mean + SD. Student’s t-test. *P < 0.05, **P < 0.01, ***P < 0.001. B) Targeted metabolomic analysis by Ultra-high performance liquid chromatography-mass spectrometry (UHPLC-MS) system. Heat map of differentially expressed metabolites in control and DCX-OE cells cultured in normal, low-glucose, glucose-deficient, and glutamine-deficient culture conditions (n=3 for each group). C) Transmission electron microscope images of Control-U251, DCX-OE, and DCX-KD cells cultured in high-glucose media, glutamine-deficient media, and glucose-deficient media supplemented with galactose. Orange arrows indicate mitochondria, white arrows indicate fusion, and black arrows indicate defective mitochondria. Scale bar, 1μm. D) Flow cytometry analysis of mitotracker red in control, DCX-OE, and DCX-KD cells. E) immunoblots of MFN1, DRP1, OPA1, and actin in Control, DCX-OE, and DCX-KD cells. F) The binding of DRP1 to FIS1, or BAX to MFN1 determined by immunoprecipitation followed by immunoblotting. G) Genset enrichment analysis for the DCX-OE versus control for GOBP_MITOCHONDRIAL_GENE_EXPRESSION. H) Seahorse metabolic assay measuring oxygen consumption rate (OCR) in Control-U251, DCX-OE, and DCX-KD cells. O, oligomycin; F, FCCP (carbonyl cyanide 4-[trifluoromethoxy] phenylhydrazone); A&R antimycin A and rotenone. I) Analysis of maximum OCR levels in normal culture conditions, no-glutamine, and galactose media. Data are mean + SD (n=3-4). Student’s t-test. *P < 0.05, **P < 0.01, ***P < 0.001. J. ECAR levels in normal culture condition, and K) analysis of maximum ECAR IN NORMAL, no-glutamine, and galactose media. Data are mean + SD (n=3-4). Student’s t-test. *P < 0.05, **P < 0.01, ***P < 0.001.

The mitochondria play crucial roles in cell functions and homeostasis; hence, mitochondria quality control is imperative for cancer cells that rely on OXPHOS. The peroxisome proliferator-activated receptor (PPAR) gamma coactivator (PGC)-1 α is a primary regulator of mitochondria biogenesis^36–38^. High DCX promoted *PGC-1α* mRNA expression (Figure S4E). Since metabolite depletion influence the mitochondria because they rely on metabolites to function, and glutamine depletion affects mitochondria morphology^39^, mitochondria integrity and biogenesis were examined via transmission electron microscopy. DCX-OE cells exhibited healthy mitochondria with defined cristae, even under glutamine-deficient conditions, whereas control and DCX-KD cells displayed swollen or fragmented mitochondria (Figure 4C). Flow cytometry analysis of mitotracker red staining further confirmed higher mitochondria activity in DCX-OE cells (Figure 4D). DCX-OE cells also exhibited higher expression of fusion proteins, MFN1 and OPA1 (Figure 4E), as we also confirmed high DRP1 and FIS1 interactions promoted by DCX overexpression for mitochondria fusion, while Bax binds to MFN1 in control and DCX-KD cells, indicating mitochondria fragmentation (Figure 4F).

GSEA demonstrated that DCX-OE cells exhibited enrichment of mitochondrial gene expression and translation which further supports enhanced mitochondrial function (Figure 4G, S4F). Next, we probed for DCX’s functions in glutamine anaplerosis and oxidative phosphorylation (OXPHOS) in real-time using Agilent’s XF Glutamine Oxidation Stress Test (which utilizes BPTES for inhibition of glutamine through GLS). In normal culture conditions, analysis of glutamine stress tests indicated significantly higher OCR in DCX-OE cells (Figure 4H, I), indicating a higher demand for glutamine for growth and metabolic activities. To study the maximal substrate uptake without bias, we grew cells in glutamine/glucose deficient culture media, thereby establishing complete depletion of extracellular and intracellular glutamine. Cellular OCR was greatly depleted across all cells (Figure 4H, I), indicating energy loss and suppressed OXPHOS due to glutamine starvation. Further analysis showed higher ECAR in DCX-OE cells across the different culture conditions, while minimal levels of ECAR were recorded in cells cultured in galactose media indicating repressed glycolysis (Figure 4J, K). Immunofluorescence imaging showed that DCX stimulated higher expression of HKII and TOM20, markers of active mitochondria, in glioma cells under both glutamine-replete and glutamine-deficient conditions (Figures S4G-L). These results are consistent with metabolomics data, further demonstrating that DCX promotes mitochondrial biogenesis.

### ATF5 and MYC regulate DCX transcription to drive GBM progression

To evaluate ATF’s role in DCX-mediated GBM progression, we inhibited ATF5 with sorafenib. Sorafenib treatment suppressed *ATF5* expression and depleted *ASNS* mRNA (Figure 5A). This is important to note since inhibition of ASNS also affected *ATF5* mRNA expression in all three types of ASNS inhibition (Figure 3), highlighting their synergistic role in DCX-driven GBM progression. Additionally, sorafenib treatment downregulated mRNA levels of *SLC7A11, BCL2, PI3K*, and *EIF2* complex genes (Figure 5A).

**Figure 5.**
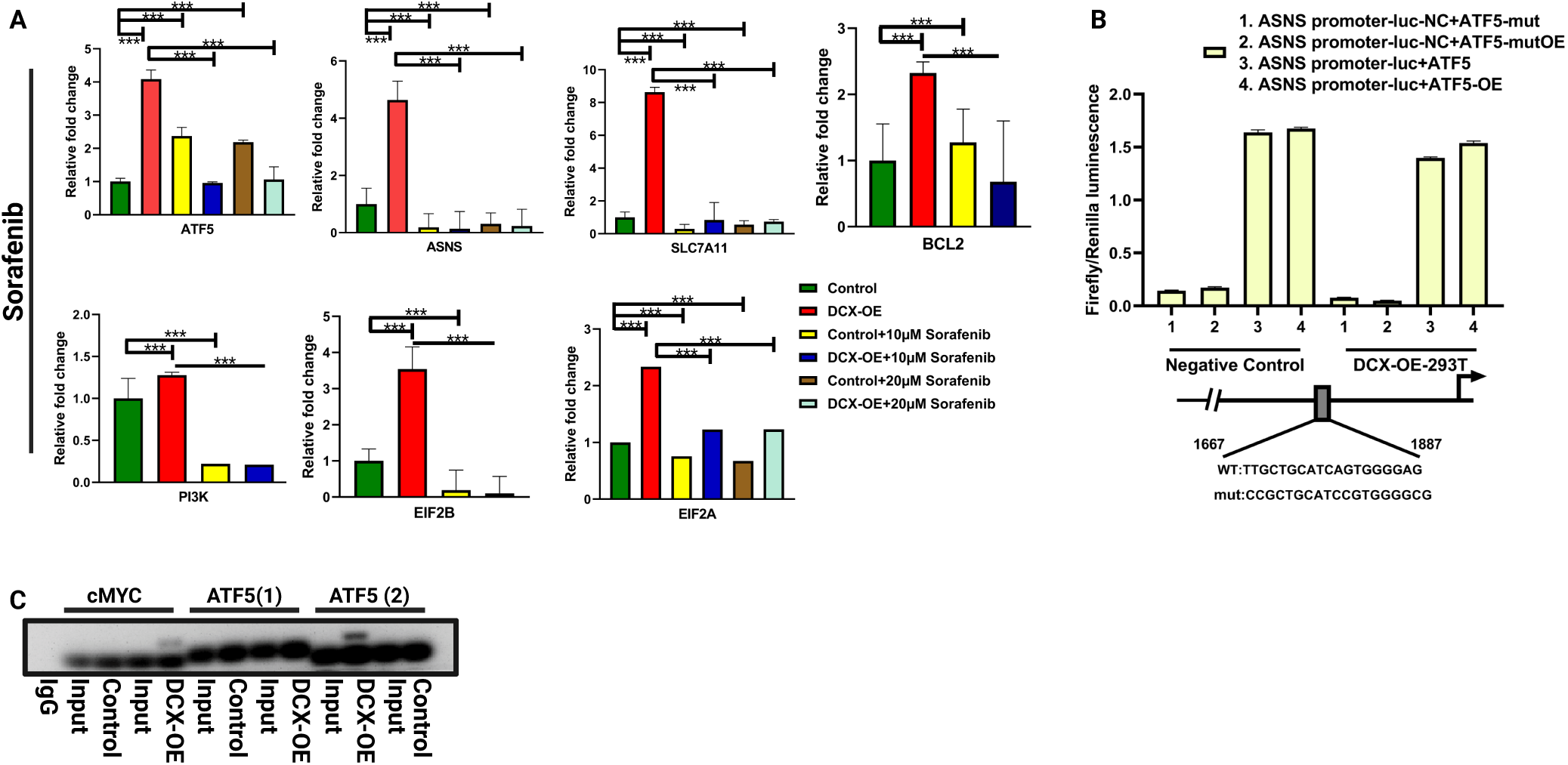
DCX regulates homeostasis via the regulation of ferroptosis in GBM cells. A) Relative fold change of mRNA levels of ATF5, ASNS, SLC7A11, PI3K, EIF2B, and EIF2A, in U251-control and DCX-OE cells treated with 10µM or 20µM sorafenib. B) Dual-luciferase Renilla/Firefly luminescence intensity of 1) ASNS promoter upstream of firefly luciferase with mutant 1 of ATF5 motif constructed into pcDNA3.1 vector, 2) ASNS promoter upstream of firefly luciferase with mutant 2 of ATF5 motif constructed into pcDNA3.1 vector, 3) ASNS promoter upstream of firefly luciferase with wild-type ATF5 motif constructed into pcDNA3.1 vector, and 4) ASNS promoter upstream of firefly luciferase with overexpression of ATF5 motif constructed into pcDNA3.1 vector. C. Semi-quantitative PCR result of Chromatin-Immunoprecipitation experiment showing the binding intensity of DCX to MYC and ATF5. ATF (1)-motif1, ATF5 (2)-motif 2. Results are presented as mean ± SEM. *P < 0.05, **P < 0.01, and ***P < 0.001.

ATF4 and ATF5 are both members of the CREB/ATF transcription factor family, and they share relatively similar sequence homology and may be functionally redundant in cancer cells^40^. Furthermore, the key features of ATF5 translational control share those described for ATF4^41,42^. Since direct promoter interactions between ATF5 and ASNS have not been previously reported, we modeled our predicted sites with already existing ATF4 binding sites on the promoter region of ASNS in 293T cells. Renilla/luminescence assay revealed luciferase activity of ATF motif on the promoter of ASNS, and mutation of the ATF motif on two different sites resulted in the inhibition of ATF5 luciferase activity on the ASNS promoter, and this was confirmed in DCX-OE glioma cells (Figure 5B). Our result confirmed that indeed, transcription factor ATF5 binds to the promoter region of ASNS at 1056 bp. Furthermore, we also sought to clarify if there is a direct link between DCX and specific transcription factors. Because the dependency on glutamine is enhanced by the expression of MYC, which drives a transcriptional program to coordinate the expression of genes involved in glutamine metabolism, and because DCX modulates the expression of ATF5 in GBM cells, we probed for cMYC-DCX and ATF5-DCX interactions to determine if these transcription factors are possible transcriptional regulator of DCX. We predicted possible binding sites on the DCX promoter region and designed primers of the DCX promoter region for chromatin-immunoprecipitation (ChIP) assays and semi-qPCR to measure the interaction of these transcription factors and DCX. Our result showed direct interactions between DCX-MYC and DCX-ATF5, indicating that ATF5 and MYC are transcriptional regulators of DCX (Figure 5C).

### DCX regulates homeostasis via the regulation of ferroptosis in GBM cells

We also confirmed enrichment of membrane transport geneset (Figure 6A), which includes SLC7A11, a top 10 upregulated DEG which also mediates metabolic reprogramming in cancer cells, leading to glucose and glutamine-dependency in cancer cells. SLC7A11 facilitates cystine import for glutathione biosynthesis and antioxidant defense to promote tumor growth partly via the suppression of ferroptosis^43^. To confirm the homeostatic effect of amino acid exchange via the high expression of xCT in DCX-OE cells and the regulation of glutamate exchange and the influx of essential amino acid coupled with the efflux of glutamate to avoid an unwanted accumulation of glutamate and maintain homeostasis, since there is a high level of glutamine synthesis and hydrolysis via ASNS and GLS in DCX-OE cells, we inhibited xCT with Erastin, a cell-permeable piperazinyl-quinazolinone compound capable of initiating ferroptosis by inhibiting cysteine glutamate antiporter system (xc-). xCT inhibition with erastin resulted in suppressed expression of ASNS, ATF5, and PI3K mRNA levels in DCX-OE cells. However, there was significantly higher *EIF2α* expression in DCX-OE cells, which may be due to an imbalanced exchange of amino acids and accumulation of excess intracellular ASNS (Figure 6B). Furthermore, *EIF2β* mRNA expression was significantly suppressed by erastin more than in the control group. Whereas, there was still higher *BCL2* mRNA expression in DCX-OE cells. However, treatment of cells with a higher dosage of Erastin (1μM) conferred total loss of these genes in both control and DCX-OE cells.

**Figure 6.**
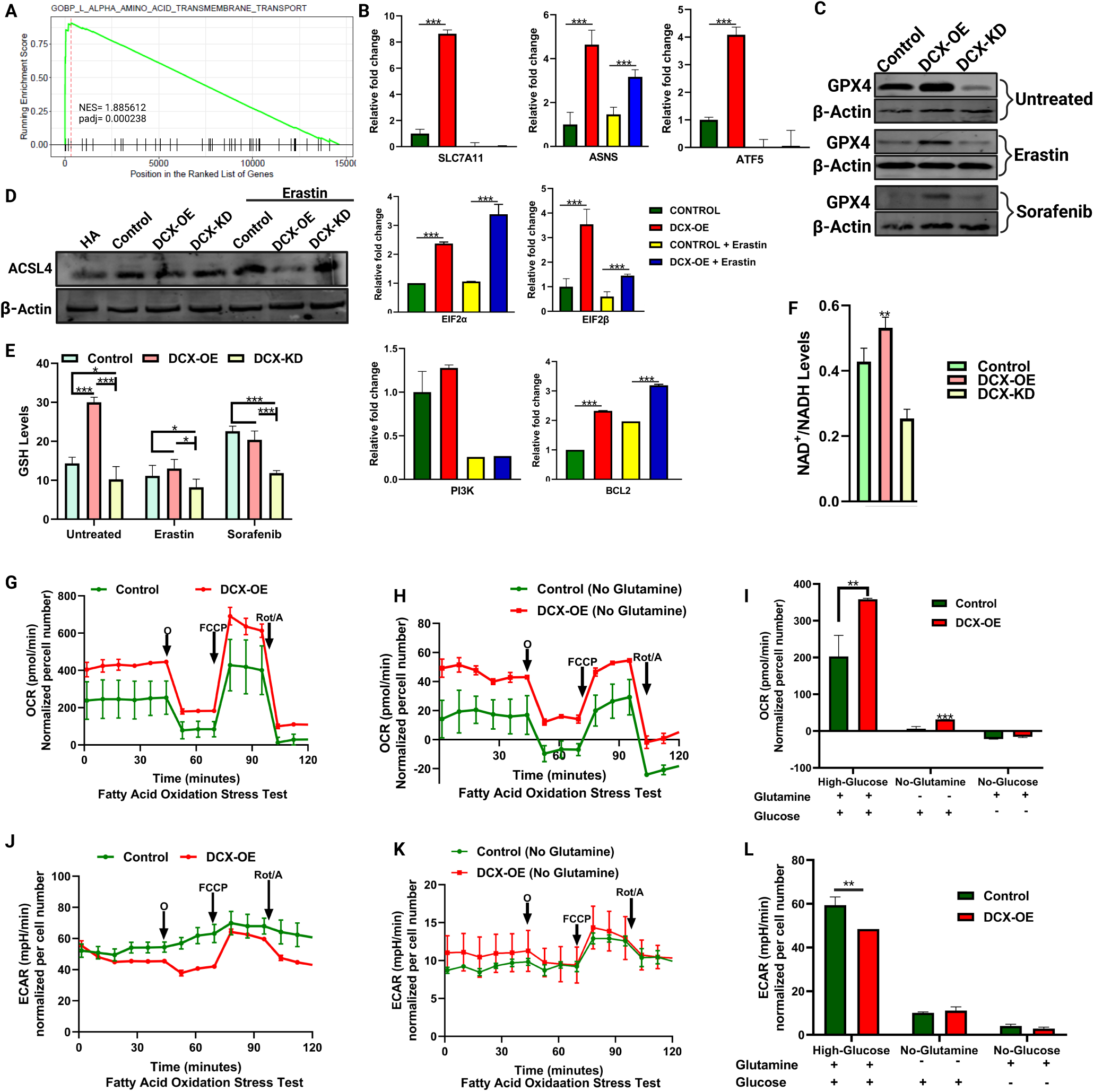
DCX regulates homeostasis via the regulation of ferroptosis in GBM cells. A) GOBP_L_ALPHA_AMINO_ACID_TRANSMEMBRANE_TRANSPORT gene set enrichment in DCX-OE cells compared with controls. B) Relative fold change of mRNA levels of SLC7A11, ASNS, ATF5, PI3K, EIF2B, and EIF2A, in U251-control and DCX-OE cells treated with erastin. C) immunoblots showing GPX4 and actin expression in cells treated with either DMSO, erastin, or sorafenib. D) Immunoblots showing ACSL4 and β-actin expression in human astrocytes (HA), U251 control, DCX-OE, and DCX-KD either untreated or treated with erastin. E) Glutathione (GSH) levels in untreated, erastin-treated, sorafenib-treated control, DCX-OE, and DCX-KD cells. F) NAD^+^/NADH levels in control, DCX-OE, and DCX-KD cells. G) Seahorse metabolic assay measuring oxygen consumption rate (OCR) in Control-U251, DCX-OE, and DCX-KD cells cultured in normal culture condition, G) or in glutamine-deficient culture condition. I) Analysis of maximum OCR levels in normal culture conditions, no-glutamine, and galactose media. J) Seahorse metabolic assay measuring extracellular acidification rate (ECAR) in Control-U251, DCX-OE, and DCX-KD cells cultured in normal culture condition, K) or in glutamine-deficient culture condition. L) Analysis of maximum ECAR levels in normal culture conditions, no-glutamine, and galactose media. Data are mean + SD (n=3-4). Student’s t-test. *P < 0.05, **P < 0.01, ***P < 0.001. O, oligomycin; F, FCCP (carbonyl cyanide 4-[trifluoromethoxy] phenylhydrazone); A&R antimycin A and rotenone. Data are mean + SD (n=3-4). Student’s t-test. *P < 0.05, **P < 0.01, ***P < 0.001.

To understand the transient reduction in ASNS and sustained globular protein synthesis in lower-dose sorafenib-treated GBM cells, we probed for the expression of glutathione peroxidase (GPX4). GPX4, a key regulator of ferroptosis is highly expressed in DCX-OE cells, which may be due to SLC7A11 stimulation. Also, treatment of cells with erastin or sorafenib significantly downregulated GPX4 expression albeit relatively higher GPX4 expression in DCX-OE cells (Figure 6C). To understand the selective inhibition of GPX4 in DCX-KD cells, we checked for ACSL4 expression. Ferroptosis induced by the inactivation of SLC7A11 or GPX4 can be abolished by the suppression of ASCL4 ^44^. Our result showed reduced ASCL4 expression in DCX-OE cells, and DCX overexpression cells react to erastin treatment by further suppression of ASCL4 expression (Figure 6D). Consistently, DCX expression correlated with GSH levels (Figure 6E). Higher total NAD^+^/NADH levels was also recorded in DCX-OE cells (Figure 6F) indicating higher redox balance in DCX-OE cells. In Long Chain Fatty Acid (LCFAs) oxidative stress tests when Etomoxir replaced BPTES or UK5099, DCX-OE cells exhibited higher OCR in normal culture conditions and glutamine-deficient culture conditions (Figures 6G-I). However, lower ECAR levels were recorded in all culture conditions (Figures 6J-L).

## Discussion

Gliomas, and GBM in particular, are notorious for being incurable in most patients ^45^, remaining a formidable clinical challenge due to their metabolic plasticity, which sustains tumor growth and contributes to therapeutic resistance ^46^. Despite advances in targeted therapy, metabolic reprogramming within tumor cells frequently enables survival under therapeutic stress. Emerging paradigm suggests that oncogenes can actively reprogram cellular metabolism to enable tumor survival and growth, redirecting metabolic pathways toward alternative energy sources, including those controlled by MAPs ^8^.

While the role of DCX in reprogrammed metabolism in cancer is largely unexplored, its crucial roles in adult neurogenesis require complete metabolic reprogramming and switching energy sources ^47^. We have previously reported that DCX promotes glioma progression, and its roles in neurogenesis make it susceptible to pro-oncogenic entities ^8,10,15^, while its knockdown leads to translocation of BAX and apoptosis via Rho-A/Net-1/p38-MAPK signaling ^48^. Our result presents an essential survival pathway for GBM cells targeting metabolism from glutamine synthesis with the aid of ASNS in glioma cells characterized by high DCX expression, coupled with the redox balance initiated by the efflux of glutamate by SLC7A11, and possible exchange for amino acids, with subsequent activation of ATF5. This study also presents DCX as a metabolic accessory, however, DCX is a neuronal MAP crucial for adult neurogenesis, a class of molecules for which the development of small-molecule drugs has proven to be challenging. Of note, the reliance of DCX-overexpressing GBM cells on glutamine metabolism highlights a vulnerability that could be targeted with glutamine antagonists, some of which are currently in clinical trials (NCT04471415) ^49^.

DCX-driven metabolic reprogramming facilitates the synthesis of essential amino acids, as demonstrated by UHPLC analysis, even under nutrient-stressed conditions. Upregulation of ASNS enables glutamine-glutamate conversion, sustaining the TCA cycle and energy generation. Glutamine is metabolized in the mitochondria via glutaminolysis, an enzymatic process involving glutamine conversion to α-ketoglutarate (αKG), a TCA cycle intermediate ^50^. Glutamine transported into the cells via the xCT is converted to glutamate and further to αKG by glutaminase (GLS). Similarly, in an ATP-dependent manner, ASNS also catalyzes the conversion of aspartate and glutamine to asparagine and glutamate in cancer cells ^51,52^.

Redox balance, the ratio between oxidizing and reducing species, is involved in regulating various signaling pathways, including the activity of protein kinases and phosphatases, which through the modulation of the posttranslational modifications of certain transcription factors regulates gene expression. Elevated SLC7A11 expression indicates controlled efflux of glutamate coupled with the inflow of cystine (to be reduced to cysteine) crucial for glutathione (GSH) biosynthesis and rapid tumor proliferation. Inhibiting SLC7A11 results in toxic glutamate accumulation and subsequent damage of macromolecules, and necrotic cell death^32,53^. Excessive ROS production or defective activation of antioxidant defenses results in oxidative damage and pathological conditions. ROS not only damages DNA, proteins, and fatty acids by direct oxidation but also activates specific signal transduction molecules that affect cell survival ^54–56^. DCX-OE cells exhibit elevated ROS and GSH levels, maintaining redox balance and survival under oxidative stress. Elevated GSH derived from ATP-dependent synthesis of glutamate, cysteine, and glycine, prevents oxidative damage and supports OXPHOS ^57^. The metabolic adaptability of DCX-OE cells ensures continued energy production under stress, with central carbon metabolomics revealing high glycine and glutamate levels, contributing to GSH abundance and homeostasis. This balance is further sustained by ASNS activity, which supports OXPHOS even as glutamate levels decline, while control cells shift towards glycolysis.

DCX also regulates OXPHOS by maintaining NADH (increased NADH-enzyme-bound fraction) and FAD (diminished FAD enzyme-bound fraction), essential co-factors for catabolic reactions of amino acid and fatty acid oxidation, glycolysis, the TCA cycle, and electron transport chain (ETC) ^58^. This ensures sustained energy replenishment in DCX-OE cells during nutrient stress. Furthermore, DCX-driven redox homeostasis enhances resistance to ferroptosis, a regulated form of cell death triggered by oxidative stress. By upregulating SLC7A11, DCX facilitates cystine import and GSH synthesis, neutralizing ROS and preventing lipid peroxidation. Combining ferroptosis inducers (e.g., erastin or RSL3) with glutamine antagonists could provide a promising therapeutic strategy to overcome metabolic plasticity in GBM.

Our findings also establish a novel relationship between DCX and ATF5, a pro-survival transcription factor involved in GBM metabolism. ATF5, highly expressed in malignant gliomas, but significantly lower in mature neurons ^59^, promotes de-novo protein synthesis in cancer cells and plays a critical role in cellular adaptation to nutrient stress by responding to amino acid limitation ^60^. Preferential targeting of cancer cells even within the limited population of ATF5-positive non-neoplastic cells is essential for high therapeutic index designed for any ATF5-targeted therapeutic intervention ^60^. DCX stimulates ATF5 expression, contributing to the anti-apoptotic features of cells exhibiting high DCX expression. Inhibiting ATF5 abrogates PI3K signaling and suppresses metabolic activity, highlighting its role as a key effector of DCX-driven metabolism. Silencing or inhibition of ASNS also represses ATF5 expression in DCX-OE cells, because ATF5 responds to amino acid limitation and promotes cellular adaptation to nutrient stress ^60,61^. Furthermore, ATF5 acts as a transcription factor for ASNS, creating a synergistic link between amino acid metabolism and cell survival. Interestingly, ATF5 and MYC also regulate DCX transcription, integrating metabolic and oncogenic signaling pathways in GBM.

Evidence in our study further supports that the interplay between ATF5, SLC7A11, and ASNS is tightly coupled with ER stress signaling, particularly through the integrated stress response involving ATF4. Prior studies have demonstrated that ASNS transcriptional activation during ER stress is primarily mediated by the PERK–eIF2α–ATF4 signaling axis, with no significant contribution from the ATF6 or IRE1/XBP1 pathways ^62^. Consistent with this, our findings show that ATF5, closely related to ATF4 acts downstream of this axis and binds directly to the ASNS promoter, positioning it as a potential co-regulator or compensatory factor in cases where ATF4 is activated. Inhibition of ASNS significantly suppresses ATF5 expression, and vice versa, suggesting a feedback loop critical for maintaining glutamine metabolism and redox homeostasis. Furthermore, the regulation of SLC7A11 by DCX and its suppression upon inhibition of ATF5 or ASNS points to a coordinated stress adaptation network sustained by ATF-family transcription factors.

In conclusion, DCX is a critical regulator of GBM metabolism, linking glutamine anaplerosis, amino acid synthesis, and redox balance to tumor survival and growth. While directly targeting DCX poses challenges dues to its role in neurogenesis, its downstream effectors-ASNS, SLC7A11, and ATF5, offer viable therapeutic targets. These findings highlight the metabolic vulnerabilities of GBM and provide a foundation for developing novel targeted therapies to exploit these weaknesses. Further research is warranted to translate these insights into effective clinical interventions.

## Ethics Statement

This study adhered to ethical standards for research involving animal subjects. All experimental procedures involving animals were conducted following approval from the Animal Care Committee of Xuzhou Medical University under protocol 202212S001.

## Author contributions

Conceptualization: AAA

Investigation: AAA, HXL, and HZ.

Writing-original draft: AAA.

Visualization: AAA, HXL, MC, BC, AIA, MMA, and AQR.

Validation: AAA.

Project administration: AAA.

Methodology: AAA, HXL, BC, HZ, KK, NI, CT, PAK, and JD.

Formal analysis: AAA, HXL, and MC.

Writing-review and editing: AAA.

Resources: AAA and ZBL.

Supervision: AAA.

Funding acquisition: ZBL

## Data Availability

All data supporting the conclusions of this study are included in the paper and Supplementary Materials. Additional data may be obtained from Dr. Ayanlaja and Xuzhou Medical University upon scientific review and completion of a material transfer agreement.

## Acknowledgments

This work is dedicated to Late Professor Dianshuai Gao, whose mentorship and funding steered the project till his demise. May his soul rest in peace! We would also like to express our gratitude to the staff of the Public Experimental Research Center of Xuzhou Medical University for allowing us to use the laboratories to complete some of our research.

