## Supplementary figures and images for "DCX Enhances Glioblastoma Metabolism Through Synergistic Regulation of Glutamine Synthesis and Metabolism-Related Genes for Cellular Homeostasis"

### Supplemental Figure 2

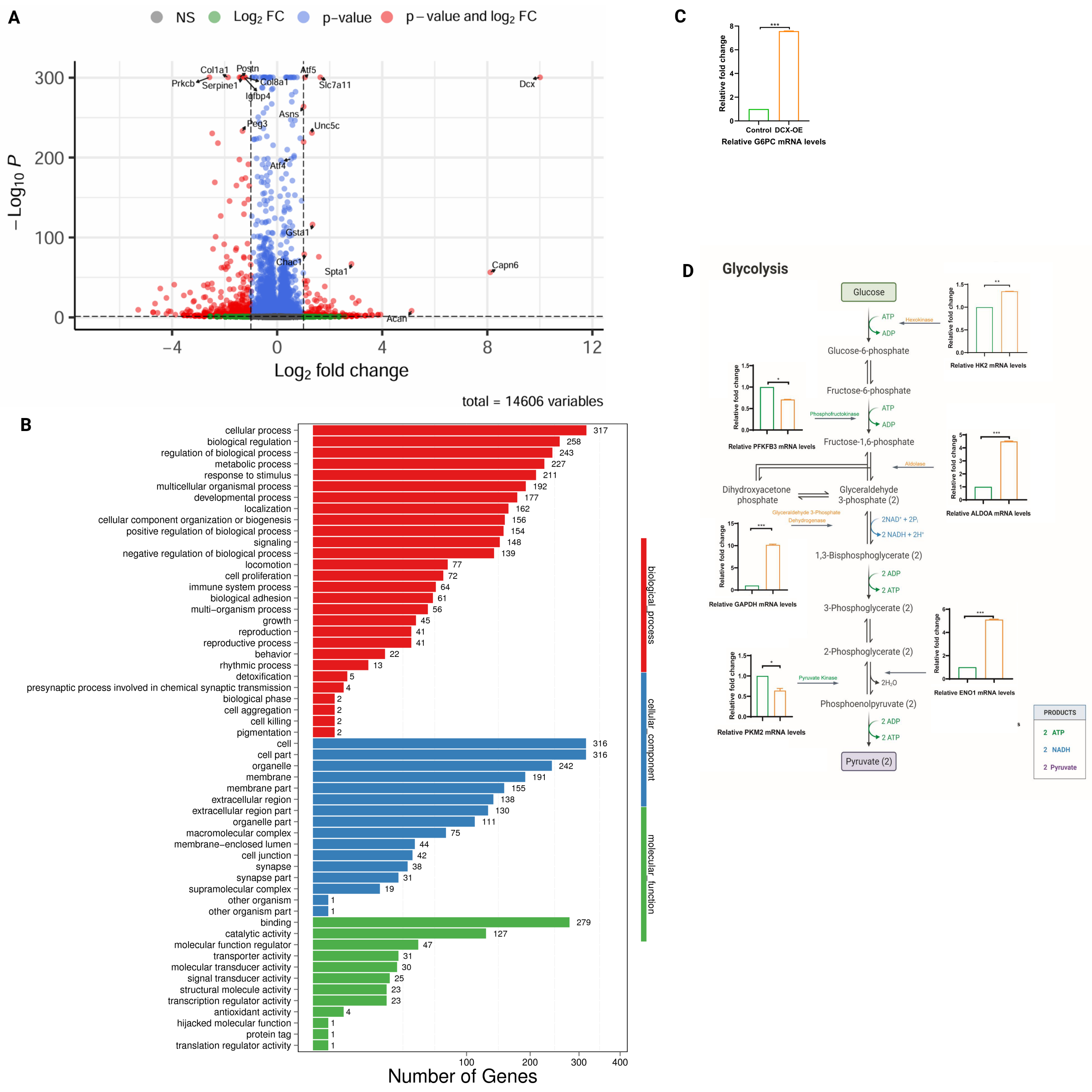

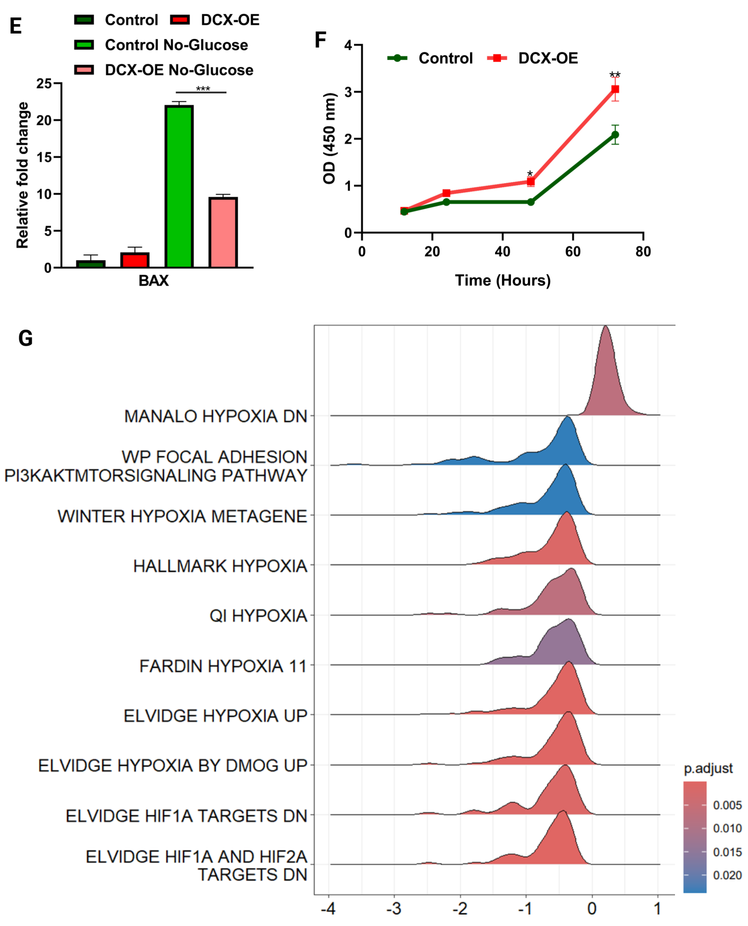

### Supplemental Figure 3

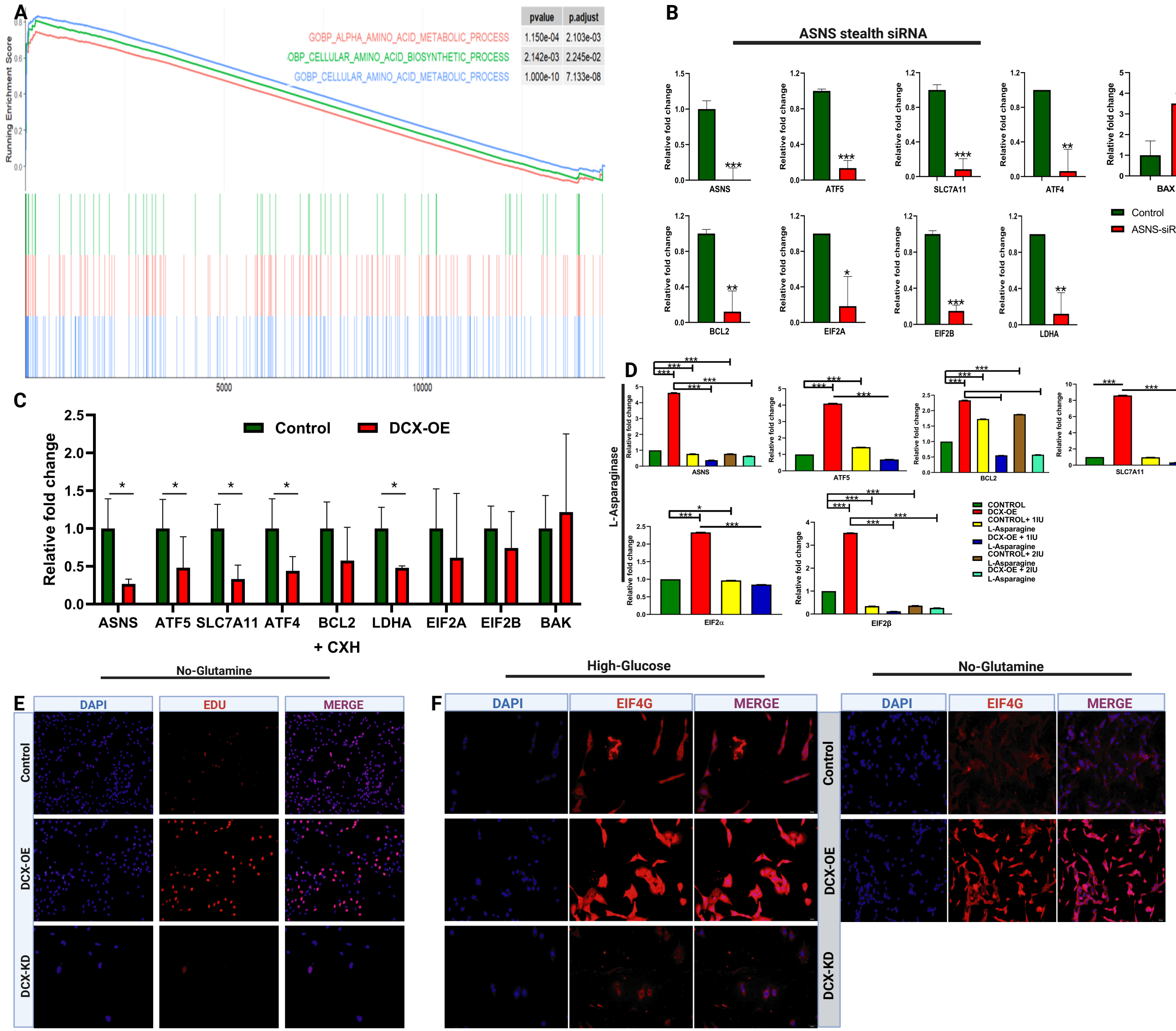

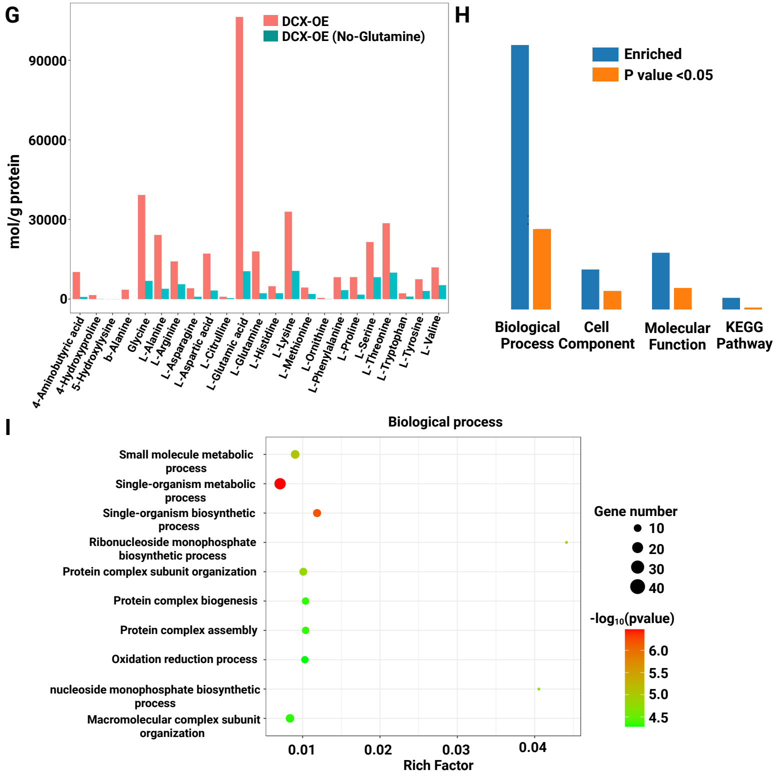

### Supplemental Figure 4

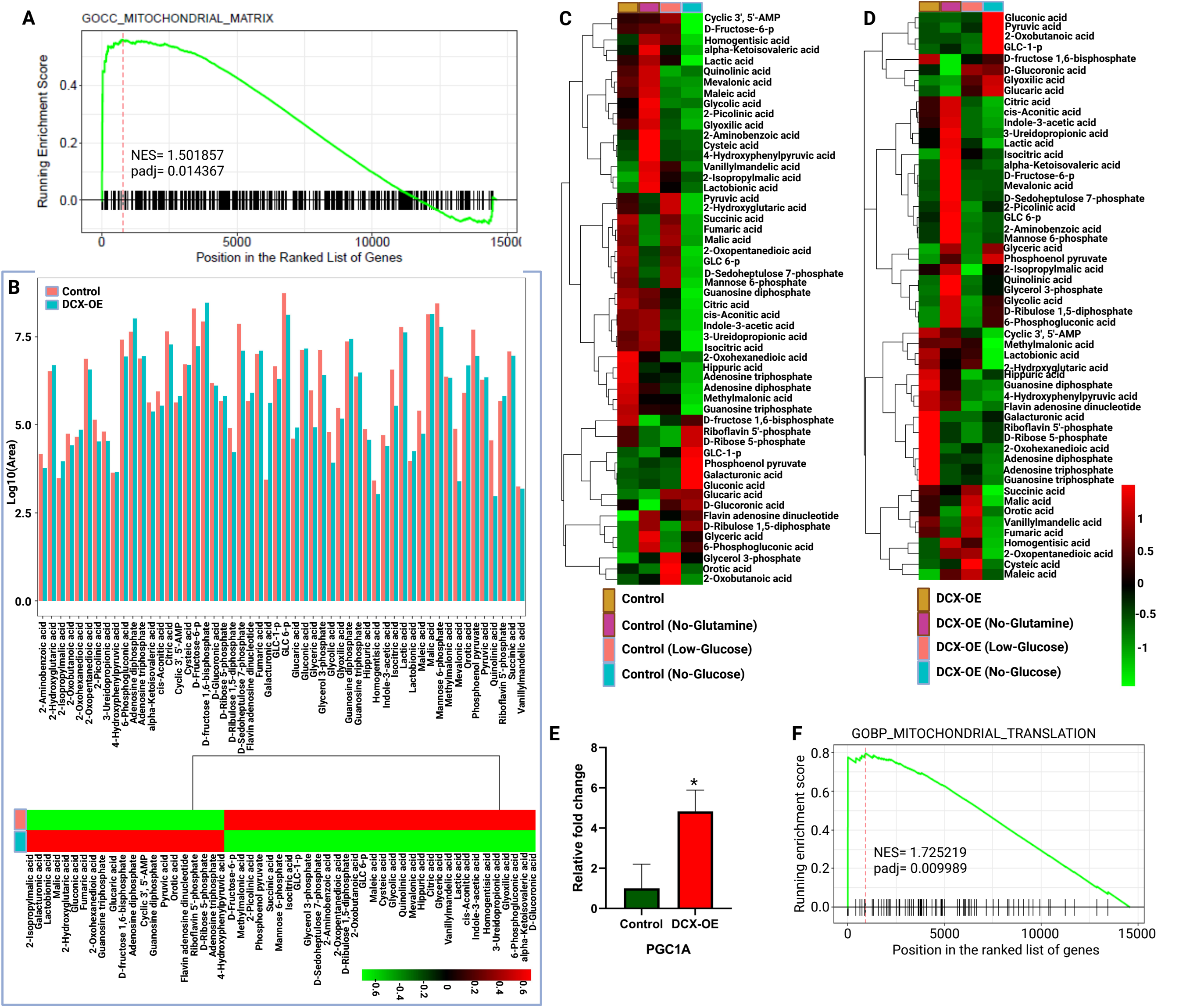

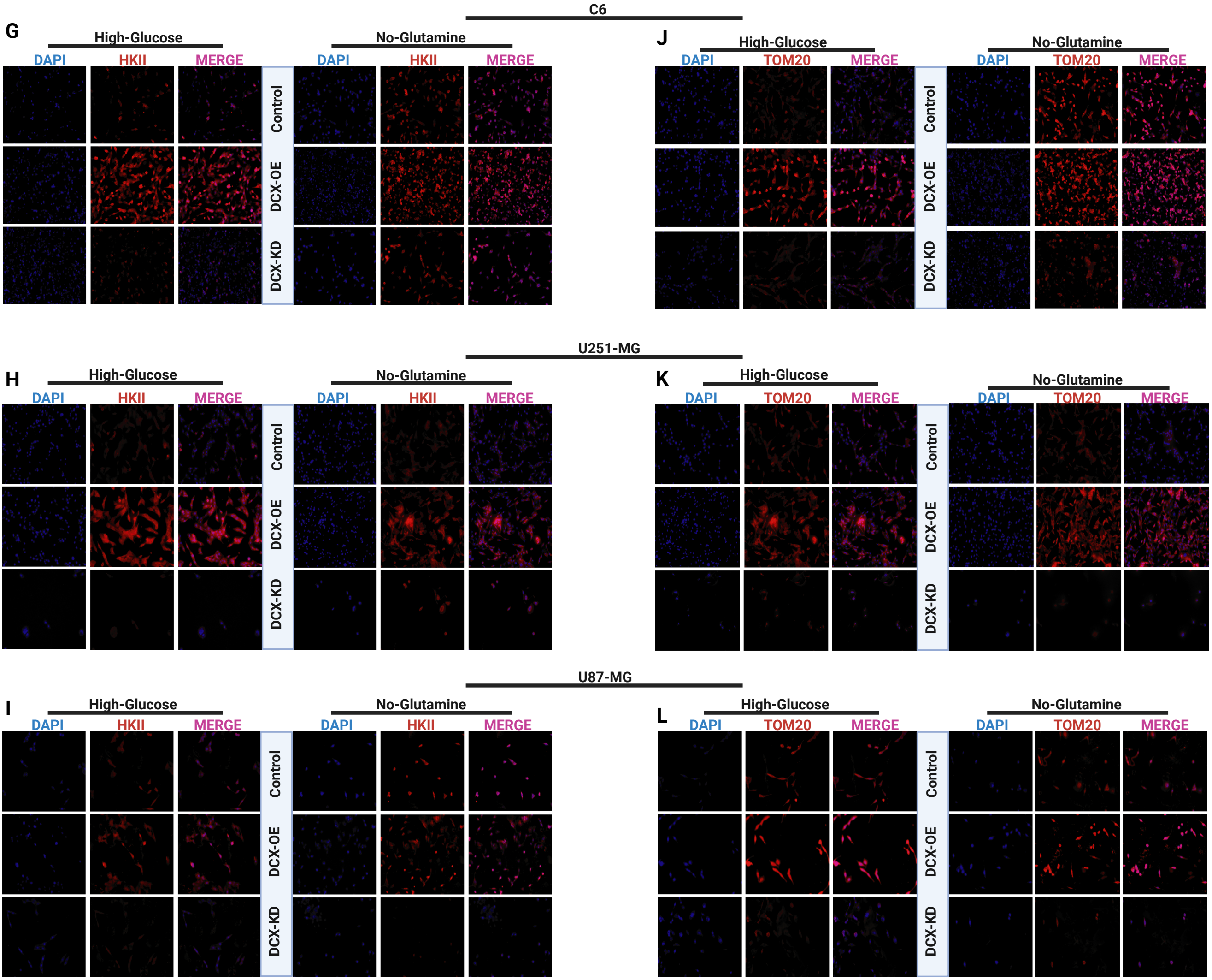

### Supplementary figures 1

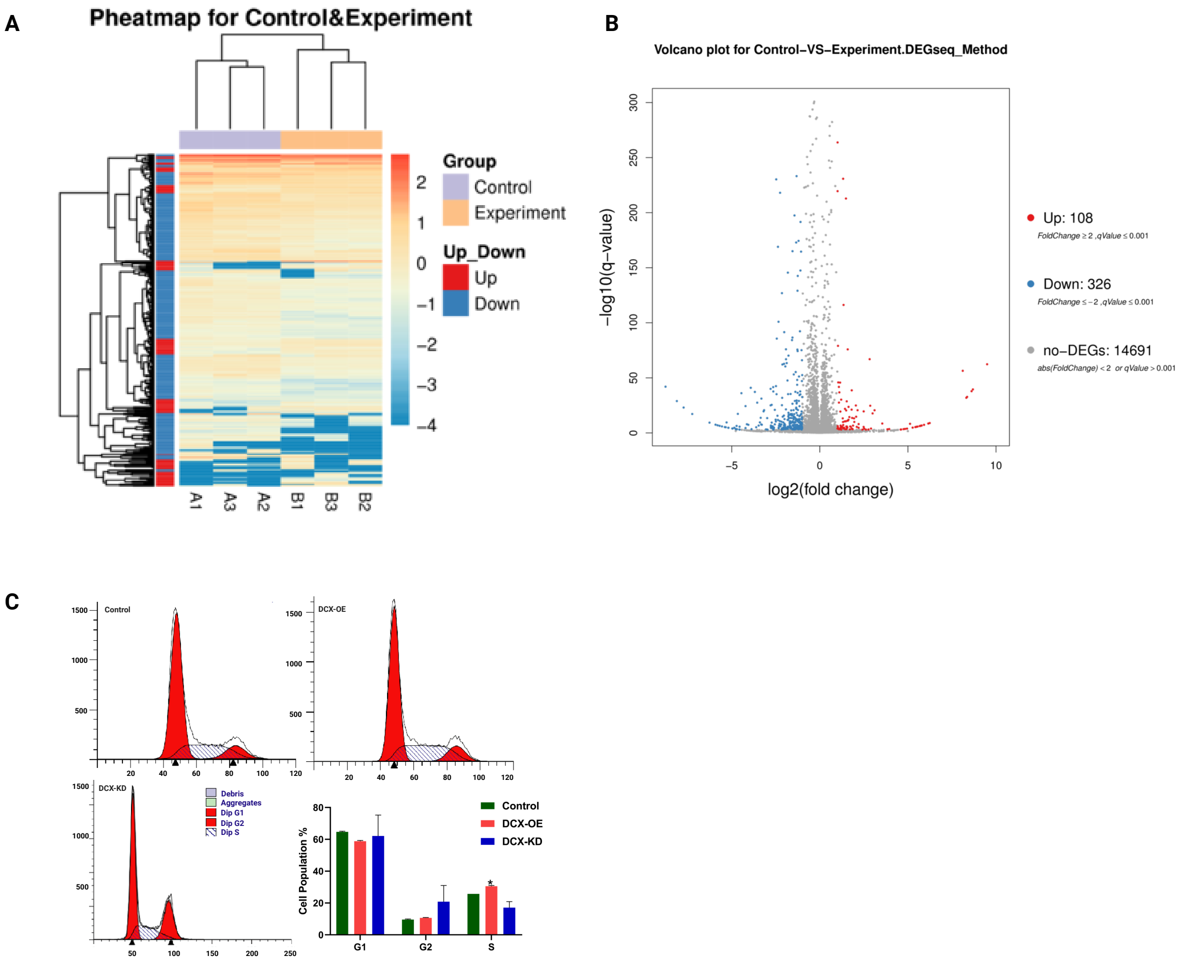
