## Supplemental File for "DCX Enhances Glioblastoma Metabolism Through Synergistic Regulation of Glutamine Synthesis and Metabolism-Related Genes for Cellular Homeostasis"

Supplementary Materials and Methods

**Antibody and other reagents information**:

| Antibody | Cat. number | company |
| --- | --- | --- |
| AKT | #9272 | Cell signaling technology |
| pAKT^s473^ | #9271 | Cell signaling technology |
| pAKT^t308^ | #9275 | Cell signaling technology |
| mTOR | ab2732 | Abcam |
| β-Actin | 66009-1-AP | Proteintech |
| pS6 | #2211 | Cell signaling technology |
| EIF4G | 15704-1-Ap | Proteintech |
| SLC7A11 | #98051 | Cell signaling technology |
| ASNS | 14681-1-Ap | Proteintech |
| ATF5 | Sc-377168 | Santacruz |
| DCX | ab18723, 4604S | Abcam, Cell signaling technology |
| PDH | ZRB2163 | Sigma |
| Vimentin | #5741 | Cell signaling technology |
| MFN1 | # 9482 | Cell signaling technology |
| DRP1 | # 8570 | Cell signaling technology |
| OPA1 | 80471 | Cell signaling technology |
| FIS1 | #32525 | Cell signaling technology |
| GPX4 | # 59735 | Cell signaling technology |
| ACSL4 | #38493 | Cell signaling technology |
| HKII | EPR20839 | Abcam |
| TOM20 | EPR15581-54 | Abcam |
| secondary antibodies | IRDye 800CW, 926-32210, | 926-32211 |
| Nestin | 19484-1-AP | Proteintech |
| secondary antibodies | Alexa fluor 488, AB_2889374, Alexa fluor 594, SA00006-8 | Proteintech |
| Hoechst | ab228550 | abcam |
| DAPI | C1002 | Beyotime |
| bovine serum albumin |  | Sigma-Aldrich |
| hematoxylin solution |  | Beyotime |
| PVDF membranes |  | Millipore |
| protein extraction reagents |  | beyotime |
| fetal bovine serum |  | gibco |
| no glucose, no glutamine, no sodium pyruvate, no phenol medium | A14430-01 | gibco |
| U87 | (RRID: CVCL_0022) | ATCC |
| U251 | (RRID: CVCL_0021) | ATCC |
| LN229 | (RRID: CVCL_0393) | ATCC |
| C6 | (RRID: CVCL_0194) | ATCC |
| Sorafenib | HY-17364 | MedChem Express |
| Erastin | HY-15763 | MedChem Express |
| L-Asparaginase | HY-P1923 | MedChem Express |
| Cycloheximide | SC0353 | Beyotime |

RT-qPCR Primers

| **Gene name** | **Abbreviation** | **Forward Primer** | **Reverse Primer** |
| --- | --- | --- | --- |
| 6-Phosphofructo-2-Kinase/Fructose-2,6-Biphosphatase 3 | PFKFB3 | AGCTTGTGCCAAAGGTCACT | AGGAGGATCTCAGGGCTCAC |
| Activating transcription factor 4 | ATF4 | ATGCCAGATGAG CTCTTGACCAC | GTCATTGTCAGAGGGAGTGTCTTC |
| Activating transcription factor 5 | AFT5 | TCAGGTACCGCCAGAGGAAGC | CTGGCTTCGTGCCTTATACACCTC |
| Aldolase, Fructose-Bisphosphate A | ALDOA | AACTTTCCTCTGCCTAGCCC | GTACAGGCACAGTCGCAGAG |
| Aconitase 2 | ACO2 | AAGGATCACTTGGTGCCTGA | GAAGGGCCCATTGATGTGT |
| Asparagine Synthetase | ASNS | TGCTTACGCCCAGATTTTCT | AAAACGGAATGCATCTGGAC |
| BCL2 Antagonist/Killer | BAK | CAGCAACATGCACAGCCTAT | CTGTGCATGTTGCTGCTGG |
| Bax | Bax | GGTTGTCGCCCTTTTCTA | CGGAGGAAGTCCAATGTC |
| Bcl-2 (B-cell lymphoma 2) | BCL2 | CATGTGTGTGGAGAGCGTCAA | GCCGGTTCAGGTACTCAGTCA |
| Beta Actin | β actin | TTGTTACAGGAAGTCCCTTGCC | ATGCTATCACCTCCCCTGTGTG |
| cystine transporter solute carrier family 7 member 11 | SLC7A11/xCT | TTTACCTGGGGATGATTTTCGA | GTGTAGAGATGAATGAAACCGC |
| Doublecortin | DCX | ATTGATGGATCCAGGAAGATCG | TGGGATTGACATTCTTGGTGTA |
| Enolase 1 | ENO1 | GCCGTGAACGAGAAGTCCTG | ACGCCTGAAGAGACTCGGT |
| Eukaryotic translation initiation factor 2A | EIF2α (EIF2A) | CTCCTGAAAGCAGCAACCTC | GACCGAGATGAAGCATCGTG |
| Eukaryotic initiation factor 2B | EIF2β (eIF2B) | GAGGCCTGAGCGAGGATTTC | CTCCGTTGTGCCTTCCAGTT |
| Fructose-1,6-bisphosphatase | FBP1 | CACAGCAGTCAAAGCCATCTCTTC | TGTTCATAACCAGGTCGTTGGAG |
| Glyceraldehyde Phosphate Dehydrogenase | GAPDH | CTGGGCTACACTGAGCACC | AAGTGGTCGTTGAGGGCAATG |
| Glucose-6-phosphatase, catalytic subunit | G6PC | CCAAGACTCCCAGGACTGGTTC | CCATGGCATGGCCAGAGGG |
| Glucose Transporter 1 | GLUT1 | TCGTCGGCATCCTCATCGCC | CCGGTTCTCCTCGTTGCGGT |
| Glutathione S Transferase Alpha 1 | GSTα1 | TCAATGCACGGGGCAGAATG | AACCATTGGCACTTGCTGGA |
| Hexokinase II | HKII | AGCCCTTTCTCCATCTCCTT | GCTTGCCTACTTCTTCACGG |
| Isocitrate Dehydrogenase (NAD(+) 3 Catalytic Subunit Alpha | IDH3A | CTGCTCAGTGCCGTGATG | TCCTCTGTGAAGTCTGAGCATTT |
| Lactate dehydrogenase A | LDHA | ATGGCAACTCTAAAGGATCAGC | CCAACCCCAACAACTGTAATCT |
| Oxoglutarate Dehydrogenase | OGDH | AGAGTCCCCTTCCCCTGAGC | TGTCCCCCGATGAAAGTGGTGGTG |
| Peroxisome proliferator- ctivated receptor gamma coactivator 1-alpha | PGC-1α | GTAAATCTGCGGGATGATGG | AATTGCTTGCGTCCACAAA |
| Phosphate-activated glutaminase | GLS2 | GACTTCTCTGGCCAGTTTGC | GCACATCATTCCCATGACATT |
| Phosphoenolpyruvate Carboxykinase 1 | PCK1 | ATGACAACTGCTGGTTGGCT | ACGTACATGGTGCGACCTTT |
| Phosphoinositide 3-kinase | PI3K | TTAGCTATTCCCACGCAGGA | CACAATAGTGTCTGTGACTC |
| Pyruvate Dehydrogenase Kinase 1 | PDK1 | CTGTGATACGGATCAGAAACCG | TCCACCAAACAATAAAGAGTGCT |
| Pyruvate kinase isozymes 2 | PKM2 | ATGTCGAAGCCCCATAGTGAA | TGGGTGGTGAATCAATGTCCA |
| Succinate Dehydrogenase Complex Flavoprotein Subunit A | SDHA | GCATTTGGCCTTTCTGAGGC | CTCCATGTTCCCCAGAGCAG |
| Succinate Dehydrogenase Complex Iron Sulfur Subunit B | SDHB | GGGGCCTGCAGTTCTTATG | AGGCGCTCCTCTGTGAAGT |
| Succinate-CoA Ligase GDP-Forming Subunit Beta | SUCLG2 | CCCTAATGTTGTGGGACAGC | GGAATTCTGCGTTGTCATCA |
| TP53-inducible glycolysis and apoptosis regulator | TIGAR | AATATGCTCCAGACTCATTCTTCC | AGGGCTCTTTAGAGAAAAAGCTG |

Supplementary Figure Legend

Supplementary Figure 1: Representative flow cytometry plots showing cell cycle distribution in U251-MG cells, DCX-overexpressing (DCX-OE) cells, and DCX-knockdown (DCX-KD) cells.

Supplementary Figure 2: A) Volcano plot labeling top 10 significant positive and negative LFC (log_2_fold change) transcripts in DCX-OE cells compared to controls. Thresholds for DEGs defined as Log2-Fold-change ≥ 1 OR ≤ -1 660 (vertical dashed lines) and Benjamini-Hochberg corrected p-values ≤ 0.05 (horizontal dashed line). B) GO analysis of enriched cellular process. C) Relative G6PC mRNA expression level in control and DCX-OE. D) Relative mRNA expression to actin probing glycolytic intermediates enzymes in U251-DCX-OE cells compared with U251-Control cells. HK II – Hexokinase II, PFKFB3- 6-Phosphofructo-2-Kinase/Fructose-2,6-Biphosphatase3, ALDOA- Aldolase, GAPDH- Glyceraldehyde3-Phosphate dehydrogenase, ENO1-Enolase1, PKM2- Pyruvate kinase isozymes 2. Three independent experiments. Data are mean + SD. Student’s t-test. *P < 0.05, **P < 0.01, ***P < 0.001. E) Relative BAX mRNA expression in control and DCX-OE cells cultured in normal or galactose media. F) CCK8 detection of cell growth in hypoxic conditions (1.3 % 02). G) Ridge plot showing distribution of log-fold changes vs control for genes within the same gene set. Results are presented as mean ± SEM. *P < 0.05, **P < 0.01, and ***P < 0.001.

Supplementary Figure 3: A) GSEA multi-sample running enrichment plot for amino acid metabolic pathways uniquely enriched in DCX-OE cells. Relative fold change of ASNS, ATF5, SLC7A11, ATF4, BAX, BCL2, EIF2A, EIF2B, LDHA mRNA expression in B) ASNS knockdown cells versus control, C) Cycloheximide (CXH)-treated and untreated cells, D) L-Asparaginase-treated and untreated. E) Representative images of EdU proliferation assay of control, DCX-OE, and DCX-KD cells cultured in glutamine-deficient culture media. Blue = nuclei staining for DAPI, red = EdU positive staining. Magnification, 100µm. F) Fluorescence imaging of EIF4G (red), and DAPI (blue) in U251-Control, DCX-OE, and DCX-KD cells cultured in high-glucose and glutamine-deficient conditions, 50µm. G) Intracellular amino acid profiling via UHPLC-MRM-MS/MS. Bar plots of differentially expressed amino acids in DCX-OE cells cultured in normal versus DCX-OE cells cultured in glutamine-deficient culture conditions (n=3 for each group). H) Differentially expressed binding proteins probed via mass-spectrometry for protein-protein interactions analysis of co-immunoprecipitated proteins with DCX. Blue bars depict enriched biological function, and orange bars indicate functional protein below the threshold p-value, 0.05. I) Dot plots display normalized enrichment scores and adjusted p-values for enrichment of biological processes. Results are presented as mean ± SEM. *P < 0.05, **P < 0.01, and ***P < 0.001.

Supplementary Figure 4: A) Genset enrichment analysis for the DCX-OE versus control for GOBP_MITOCHONDRIAL_MATRIX. Targeted metabolomic analysis by UHPLC-MS system B) comparing control and DCX-OE, C) control cells cultured in normal, glutamine-deficient, low-glucose, and glucose-deficient culture conditions or, D) DCX-OE cells cultured in normal, glutamine-deficient, low-glucose, and glucose-deficient culture conditions. E) Relative fold change of PGC1α mRNA expression in control and DCX-OE cells. F) Genset enrichment analysis for the DCX-OE versus control for GOBP_MITOCHONDRIAL_TRANSLATION. Fluorescence imaging of (G-I) HKII Hexokinase2, red), or (J-L) TOM20, red and DAPI (blue) in U251-Control, DCX-OE, and DCX-KD cells cultured in normal or glutamine-deficient culture condition, 50µm. Statistical analyses were conducted using an unpaired two-tailed Student’s t-test for comparisons between groups B-C and E-F. Results are presented as mean ± SEM. *P < 0.05, **P < 0.01, and ***P < 0.001.
